# Single-Cell Multi-Omics Dissects Transcript Isoform and Immune Repertoire Dynamics in Human Immunosenescence

**DOI:** 10.1101/2025.11.21.689731

**Authors:** Yuhui Zheng, Ze-Hui Ren, Yafei Yang, Wenteng Liu, Fupeng Li, Yajun Zuo, Tao Zeng, Xue Wang, Xiumei Lin, Xinyue Zhu, Baibing Guan, Wenwen Zhou, Yunting Huang, Wangsheng Li, Yu Feng, Xiao Yang, Xin Liu, Xin Jin, Hanjie Li, Yuliang Dong, Xiangliang Yang, Xun Xu, Jianhua Yin, Chuanyu Liu

## Abstract

Immunosenescence, a major hallmark of systemic aging, refers to the progressive functional decline of the immune system. This decline not only compromises host defense and immunological memory but also fuels chronic inflammation and tissue degeneration (collectively known as inflammaging). While single-cell RNA sequencing (scRNA-seq) has revealed transcriptomic alterations in immune aging, analyses restricted to transcript abundance fail to capture deeper regulatory layers, such as transcript isoform diversity and the remodeling of immune receptor repertoires. To address this, we present the human peripheral immune single-cell multi-omics atlas that integrates gene expression, transcript isoforms diversity, and immune receptor repertoires. By combining single-cell full-length transcriptome sequencing (scCycloneSEQ), short-read scRNA-seq, and single-cell immune receptor sequencing (scTCR/BCR-seq), we systematically profiled peripheral blood mononuclear cells (PBMCs) from healthy young and elderly donors. Our analyses uncovered extensive age-related remodeling of immune cell composition, functional states, and TCR/BCR diversity. Notably, we identified in CD4⁺ effector memory T cells exhibited widespread differential isoform usage (DIU), 3′UTR length variation, and a marked reshaping of cytotoxic T lymphocyte (CTL) clonotypes—all closely associated with aging-related inflammation and cellular senescence. This multi-omics atlas delineates key molecular features of immunosenescence and provides a high-resolution resource for deciphering the regulatory architecture underlying immune aging.

## INTRODUCTIONS

Aging is a multifactorial biological process characterized by a progressive decline of physiological functions, constituting the foremost risk factor for numerous chronic diseases (Kennedy et al., 2014). The immune system is particularly susceptible to age-related deterioration, a dysfunction known as immunosenescence. Recognized as a central mechanism in organismal aging, immunosenescence not only propels systemic functional decline but also serves as a critical barrier to extending health span (Borgoni et al., 2021; Liu et al., 2023; Walford, 1964). This condition is characterized by the depletion and functional attenuation of specific immune cell populations, resulting in impaired defense against pathogens and a weakened establishment of long-term immunological memory (Stahl and Brown, 2015). These deficits manifest clinically as reduced responses to infections and vaccinations, coupled with a chronic, low-grade sterile inflammatory state termed “inflammaging” (Ferrucci and Fabbri, 2018). Accumulating evidence indicates that immune aging is not an isolated phenomenon; rather, it plays a causal role by actively contributing to the functional decline of peripheral organs, thereby driving the systemic aging process (Ferrucci and Fabbri, 2018).

Conventional high-throughput transcriptomics, particularly short-read single-cell RNA sequencing (scRNA-seq), has greatly advanced our understanding of age-related transcriptional changes across diverse immune cell subsets. Yet, because it primarily quantifies gene-level expression, this approach provides limited insight into transcript isoform diversity and other regulatory layers that shape immune cell function (Harries et al., 2011; Huang et al., 2021; Mogilenko et al., 2021a; Terekhova et al., 2024; Wang et al., 2025a; Yin et al., 2025; Zhong et al., 2023). The functional state of the immune system is shaped by both regulation of transcript architecture and receptor repertoire diversity [14-16]. At the transcript architecture level, alternative splicing and differential promoter/polyadenylation site usage generate distinct isoforms that can fundamentally alter the properties of immune molecules, directly influencing processes such as antigen presentation and T cell activation thresholds (Banerjee et al., 2023). At the repertoire level, the adaptive immune system’s strength lies in the vast diversity of its T cell and B cell receptors (TCR/BCR). Aging induces repertoire contraction and clonal expansion, eroding this diversity and diminishing immune adaptability (Zhong et al., 2023). Therefore, a comprehensive dissection of immunosenescence requires moving beyond gene-level quantification to systematically map the dynamics of transcript isoforms and immune receptor repertoires.

To bridge this gap, we established a high-resolution single-cell multi-omics framework that integrates full-length transcriptome sequencing (scCycloneSEQ), short-read scRNA-seq, and single-cell immune receptor sequencing (scTCR/BCR-seq). By profiling peripheral blood mononuclear cells (PBMCs) from nine young and ten elderly healthy donors, we uncovered extensive age-related remodeling of immune cell composition, functional states, and receptor diversity. Crucially, our analysis uncovered widespread and cell type-specific shifts in transcript isoform usage, UTR variation, and alternative splicing events that are invisible to standard methods. Mechanistically, we identified distinct aging-associated functional states within CD4⁺ effector memory T cells, characterized by the activation of senescence and inflammatory programs, and linked this transcriptomic remodeling directly to the functional decline of the aging immune system. This study establishes a foundational multi-omics atlas of human immunosenescence, offering novel mechanistic insights and a valuable resource to the scientific community for developing targeted interventions to preserve immune function during aging.

## RESULTS

### Construction of a High-Resolution Omics Atlas of Peripheral Immune Cells Across Age Groups

To comprehensively profile the impact of age on the composition, function, and transcriptional regulation of peripheral immune cells, a multi-modal single-cell analysis was performed on PBMCs from ten older adults (60-70 years; five males, five females) and nine younger adults (30-40 years; four males, five females), all of whom were self-reported to be free of active disease at the time of sampling. For each cell, three complementary data types were acquired and integrated: short-read single-cell RNA sequencing (scRNA-seq) for gene expression profiles, long-read single-cell sequencing for full-length transcript isoforms, and single-cell immune receptor sequencing (scTCR/BCR-seq) for paired receptor sequences (Figure 1A).

**Figure 1.**
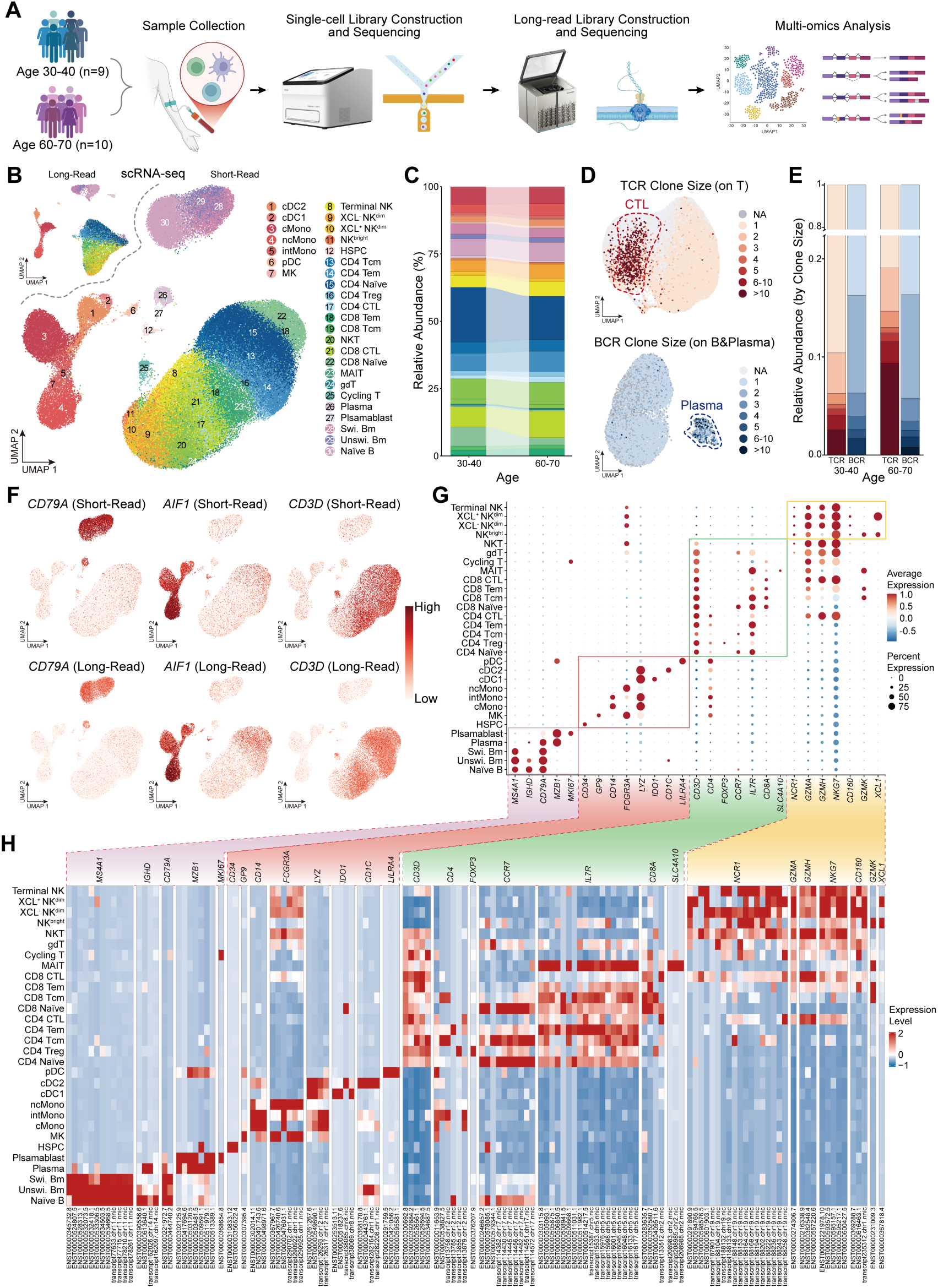
Multi-omics Integration and Cellular Landscape of Human Peripheral Immune Cells. **A**, Study design overview, illustrating the collection of peripheral blood samples and the integration of short-read scRNA-seq, long-read transcriptome sequencing, and single-cell TCR sequencing for multi-omics analysis. **B**, Uniform manifold approximation and projection (UMAP) visualization of 103,879 high-quality PBMCs retained after quality control and doublet removal. Cells are classified into 30 immune cell subtypes, each represented by a distinct color. Top left: long-read scRNA-seq; middle: short-read scRNA-seq. **C**, Proportional distribution of 30 immune cell subtypes between the two groups. Each color represents a distinct immune cell subtype. **D**, UMAP visualization of T cells showing the distribution of TCR clone sizes (top) and B cells showing the distribution of BCR clone sizes (bottom). **E**, Distribution of TCR clone sizes and BCR clone sizes between the two groups. **F**, Uniform manifold approximation and projection (UMAP) visualization showing the expression patterns of representative marker genes (*CD79A*; *AIF1*; *CD3D*) across long-read and short-read datasets. **G-H**, Expression profiles of representative marker genes across major immune cell subsets. Results derived from short-read (G) and long-read (H) data are shown separately. The color scale from blue to red indicates low to high expression levels.

Following rigorous quality control procedures and doublet removal from PBMCs (see Methods), we retained 103,879 high-quality cells from short-read scRNA-seq. For systematic annotating the immune cells phenotypes, we constructed a high-resolution cellular atlas base on scRNA-seq data. Through an iterative process of dimensionality reduction and clustering, followed by annotation based on the expression of canonical lineage markers, we classified cells into five major lineages at Level 1 (L1), including CD4 T cells, CD8 T and other T cells, B and Plasma cells, Hematopoietic Stem and Progenitor Cells (HSPCs) and Myeloid cells, and Natural Killer (NK) cells (Figure S1A and C). Both age groups display typical immune cell distributions in Uniform Manifold Approximation and Projection (UMAP) space, indicating consistent cell type definitions across cohorts (Figure S1B), with cell numbers per subject and UMAP profiles provided in Figure S2. Building on this conserved architecture, cells were further classified through subclustering informed by established marker genes, ultimately resolving 30 distinct peripheral immune cell subsets (Figure S1B and C; Figure S3; Figure S4).

Our study design enabled the simultaneous acquisition of full-length transcripts, short-read gene expression profiles, and paired immune receptor (TCR/BCR) from the same single cells by leveraging shared cell barcodes across sequencing platforms. This integrative strategy provided a comprehensive view of gene expression, transcript isoform abundance and usage, and immune receptor diversity at single-cell resolution. The final high-quality dataset comprises 103,869 single cells, capturing 30,771 expressed genes, 78,401 transcript isoforms, and fully reconstructed immune receptor repertoires through short-reads, including 11,027 paired αβ TCRs and 5,678 paired BCRs, constituting a high-resolution multi-omics resource (Figure 1B and D).

To characterize the adaptive immune receptor landscape, we constructed a global clonotype network capturing both B-cell (Figure S5A and B) and T-cell receptor repertoires (Figure 6A and B). Per-sample analysis of TCR clone size dynamics revealed that the majority of expanded clonotypes were predominantly represented in the older cohort and were concentrated within cytotoxic T lymphocyte (CTL) subsets, highlighting a key difference between the Age 30-40 and Age 60-70 cohorts (Figure 1D and E). In parallel, analysis of the B-cell receptor repertoire revealed clonal expansion, particularly within Plasma cells, although no significant differences were observed between age groups (Figure 1D and E). Additionally, gene expression profiles inferred from short- and long-read data within the same samples showed high concordance (Figure 1F; Figure S7A), and the expression patterns of canonical marker genes were highly reproducible across both sequencing platforms (Figure 1G and H). Long-read sequencing also provided uniform coverage across gene bodies, enabling the accurate detection of transcript isoforms (Figure S7B). The long-read component substantially expanded the transcriptomic landscape, identifying 17,466 novel transcripts, including novel isoforms with known splice junctions (NIC) and those containing previously unannotated splice sites (NNIC) relative to the GENCODE v38 reference (Figure S7C and D). These newly detected isoforms, which span a broad range of transcript lengths, further demonstrate the depth, comprehensiveness, and reliability of our dataset.

Overall, this study establishes a comprehensive, high-resolution dataset that constitutes a valuable resource for systematic analyses of gene expression, transcript isoform regulation, and immune receptor remodeling in aging immune cells.

### Aging Remodels the Composition and Transcriptomic Landscape of T Cells

To dissect the impact of aging on the immune system at both the cellular composition and transcriptional levels, we began by comparing the subset landscape of PBMCs between the Age 30-40 and Age 60-70 cohorts. Consistent with previous findings, we observed significant immune remodeling in the older cohort, which was characterized by a marked reduction in CD8⁺ naïve T cells and a concomitant increase in regulatory T cells (Tregs) within the CD4⁺ compartment (Figure S8) (Garg et al., 2014; Luo et al., 2022; Terekhova et al., 2023a; Wang et al., 2025b). The decline in CD8⁺ naïve T cells align with age-related thymic involution and is expected to respond to novel antigens (Lynch et al., 2009; Sauce et al., 2009). Concurrently, the expansion of Tregs, which suppress T cell activity, reflects a shift toward an immunosuppressive state associated with age-related conditions such as infections, cancer, and autoimmune disorders (Ye et al., 2012).

We next sought to identify the immune cell type whose gene expression is most affected by aging. Ranking all 30 cell types by their number of differentially expressed genes (DEGs) revealed that both T cell lineages and myeloid cells exhibited significant remodeling of its gene expression profile (Figure 2A). Enrichment analysis indicated that genes upregulated in the younger cohort were significantly associated with key immune processes, including lymphocyte differentiation, B cell receptor signaling, adaptive immune response, T cell-mediated cytotoxicity, lymphocyte-mediated immunity, and humoral immune response (Figure 2B). To further quantify the aging status of individual immune cell subsets, we computed an immunosenescence-associated gene set (IAG) score (Liu et al., 2025). This analysis demonstrated a systemic impact of aging on T cells, with significantly elevated IAG scores in the Age 60-70 cohort across all T cell subsets. The most pronounced differences were observed in multiple CD4^+^ and CD8^+^ populations (*P* < 0.01) (Figure 2C). Consistent with existing research, our findings collectively indicate an association between T cells and the immune response in aging (Terekhova et al., 2023b; Wang et al., 2025b).

**Figure 2.**
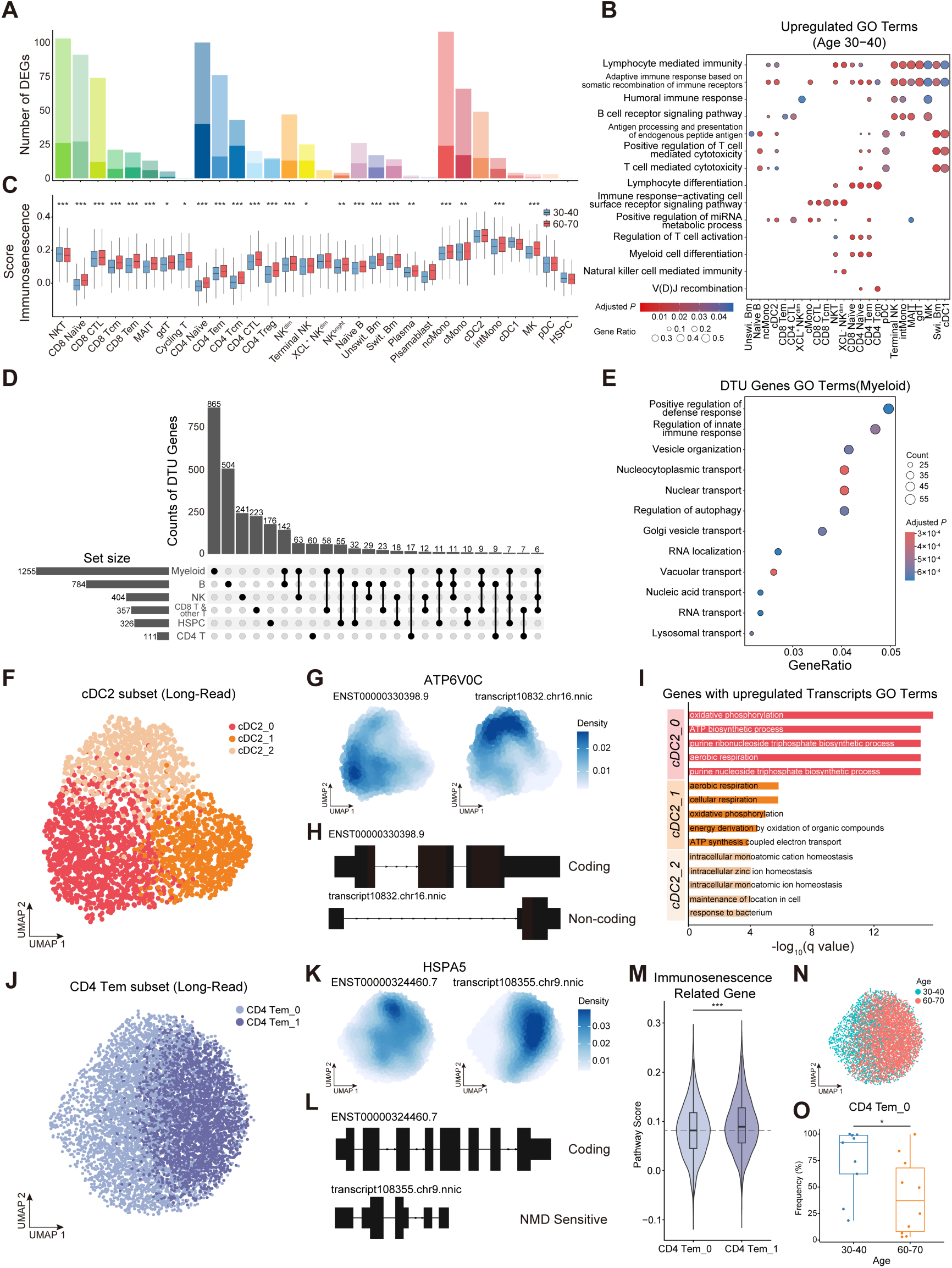
Multi-omics Profiling Reveals Aging-associated Alterations in Transcriptional and Isoform Landscapes Across T Cell and Myeloid Cell Subsets. **A**, Bar plot illustrating the count of differentially expressed genes identified in scRNA-seq across all cell subsets. **B**, Gene Ontology (GO) enrichment analysis of DEG in all cell subsets. **C**, Comparison of immunosenescence scores between groups across all cell subsets. **D**, UpSet plot illustrating the number of genes exhibiting differential transcript usage (DTU) across different cell types. **E**, Gene Ontology (GO) enrichment analysis of DTU-associated genes in the myeloid cell population. **F**, UMAP of the cDC2 population derived from long-read sequencing, with cells colored by the three cDC2 subsets to illustrate intra-type heterogeneity. **G**, The UMAP plot displays expression of two distinct ATP6V0C transcripts within the cDC2 subset. **H**, Schematics of the transcript structures for the two isoforms of ATP6V0C shown in (G). **I**, GO enrichment analysis of upregulated genes in each cDC2 subset, displaying the top enriched biological process terms. **J**, UMAP of the CD4 Tem population derived from long-read sequencing, with cells colored by the two CD4 Tem subsets. **K**, The UMAP plot displays expression of two distinct HSPA5 transcripts within the CD4 Tem subset. L, Schematics of the transcript structures for the two isoforms of HSPA5 shown in (K). **M**, Violin plot comparing the pathway scores for an immunosenescence-related gene set between the CD4 Tem_0 and CD4 Tem_1 subclusters. **N**, UMAP of CD4 Tem cells colored by donor age group (30-40 *vs.* 60-70 years) to visualize age-associated shifts in distribution. **O**, Box plot comparing the frequency of the CD4 Tem_0 subset between the 30-40 and 60-70 age groups.

### Single-Cell Isoform Analysis Reveals Aging-Associated Functional Subgroup in T Cells and Myeloid Cells

While our gene expression analyses revealed substantial differences in immune cell composition and expression between middle-aged and elderly individuals, focusing solely on total gene expression may obscure changes in transcript abundance and usage within individual genes. To address this, we performed a differential transcript usage (DTU; also referred to as DIU; Methods) analysis, focusing on transcript isoform dynamics between the Age 30-40 and Age 60-70 groups. Across all cell populations, we detected 3,944 transcripts with significant differential usage, corresponding to 2,617 genes (*P* < 0.05, |dIF (difference in Isoform Fraction)| > 0.1), with these DTU events widespread yet exhibited pronounced cell type-specific enrichment (Figure 2D). Most DTU genes in each cell type were not detected as DEGs (Figure S9A), underscoring that conventional gene-level analyses miss a substantial layer of intra-locus heterogeneity. Notably, among the various cell types, myeloid cells exhibited the highest number of DTU events (Figure 2D). Gene Ontology enrichment analysis revealed that these DTU genes were significantly associated with pathways such as “positive regulation of cytokine production”, “positive regulation of defense response,” and “regulation of innate immune response” (Figure 2E), highlighting pronounced isoform-level heterogeneity within this lineage between middle-aged and elderly individuals.

Building on the observation that myeloid cells harbor the highest number of DTU events, we next investigated whether this isoform-level heterogeneity reflects functionally distinct subpopulations within myeloid lineages. Notably, ROGUE (Ratio of Global Unshifted Entropy)(Terekhova et al., 2023b; Wang et al., 2025b) analysis of the myeloid subset cDC2 revealed relatively elevated transcriptional heterogeneity, suggesting that this population may encompass discrete functional subclusters (Figure S9B). Leveraging the high resolution of single-cell long-read sequencing, we indeed resolved cDC2 into three distinct subgroups (Figure 2F). Interestingly, differential usage of two ATP6V0C isoforms was observed across these subgroups (Figure 2G and H), with the cDC2_0 subgroup preferentially expressed the canonical full-length ATP6V0C isoform which is predicted to retain full functionality. ATP6V0C encodes a core subunit of the V-ATPase complex, a critical mediator of end lysosomal acidification required for antigen processing and presentation in cDC2s (Binnewies et al., 2019; Ten Broeke et al., 2013). The preferential use of these functionally intact isoforms suggest that cDC2_0 may specialize for efficient antigen processing and presentation. This interpretation is further supported by Gene Ontology (GO) enrichment analysis of upregulated genes, which indicated a significant enrichment of energy metabolism-related pathways within the cDC2_0 subgroup (Figure 2I).

In addition, although CD4 T cell populations exhibited relatively few DTU events, single-cell long-read sequencing enabled the identification of two transcriptionally distinct subsets within the CD4 Tem compartment (Figure 2J). These subgroups exhibited differential usage of multiple isoforms, including those of HSPA5 (Figure 2K). HSPA5, a key chaperone in the unfolded protein response (UPR), mitigates ER folding stress during TCR activation (Hetz, 2012; Thaxton et al., 2017). The CD4 Tem_1 subgroup preferentially expressed a truncated HSPA5 isoform, which may compromise ER stress handling and thereby alter functional responses upon antigen reencounter (Figure 2L). To evaluate whether transcript-level differences translate into broader aging-associated functional states, we calculated an aging-related gene set scoring approach to each CD4 Tem cell, revealing that CD4 Tem_1 exhibited significantly higher senescence scores than Tem_0 (Figure 2M), consistent with an age-like functional state. Supporting this, proportion analysis demonstrated enrichment of CD4 Tem_1 in the elderly cohort (Figure 2N and O). Collectively, these findings position CD4 Tem_1 as a functionally aged, senescence-prone memory T cell subgroup that may be a key contributor to immunosenescence.

### Single-Cell 3′UTR Profiling Reveals Post-Transcriptional Regulation Underlying Immunosenescence

To further dissect the molecular underpinnings of immunosenescence, we performed a comprehensive analysis of alternative splicing (AS) events and transcription start site (TSS) and polyadenylation site (PAS) between age groups. Aging was associated with widespread transcriptomic remodeling, including canonical AS events such as exon skipping and alternative splice site selection (Figure S10A). Beyond AS, we found that genes exhibited greater diversity in PAS usage than in TSS usage (Figure 3A and B), indicating a relative enrichment of 3′ end heterogeneity. Interestingly, genes harboring a higher number of alternative TSSs also tended to utilize a more diverse set of PASs, whereas genes with highly diverse PAS usage did not necessarily show increased TSS diversity (Figure 3C; Figure S10B). This asymmetric relationship, in which 3′ end variability is not strictly coupled to 5′ end variability, suggests a potentially greater role for 3′ end-mediated regulation in transcriptome landscape during aging.

**Figure 3.**
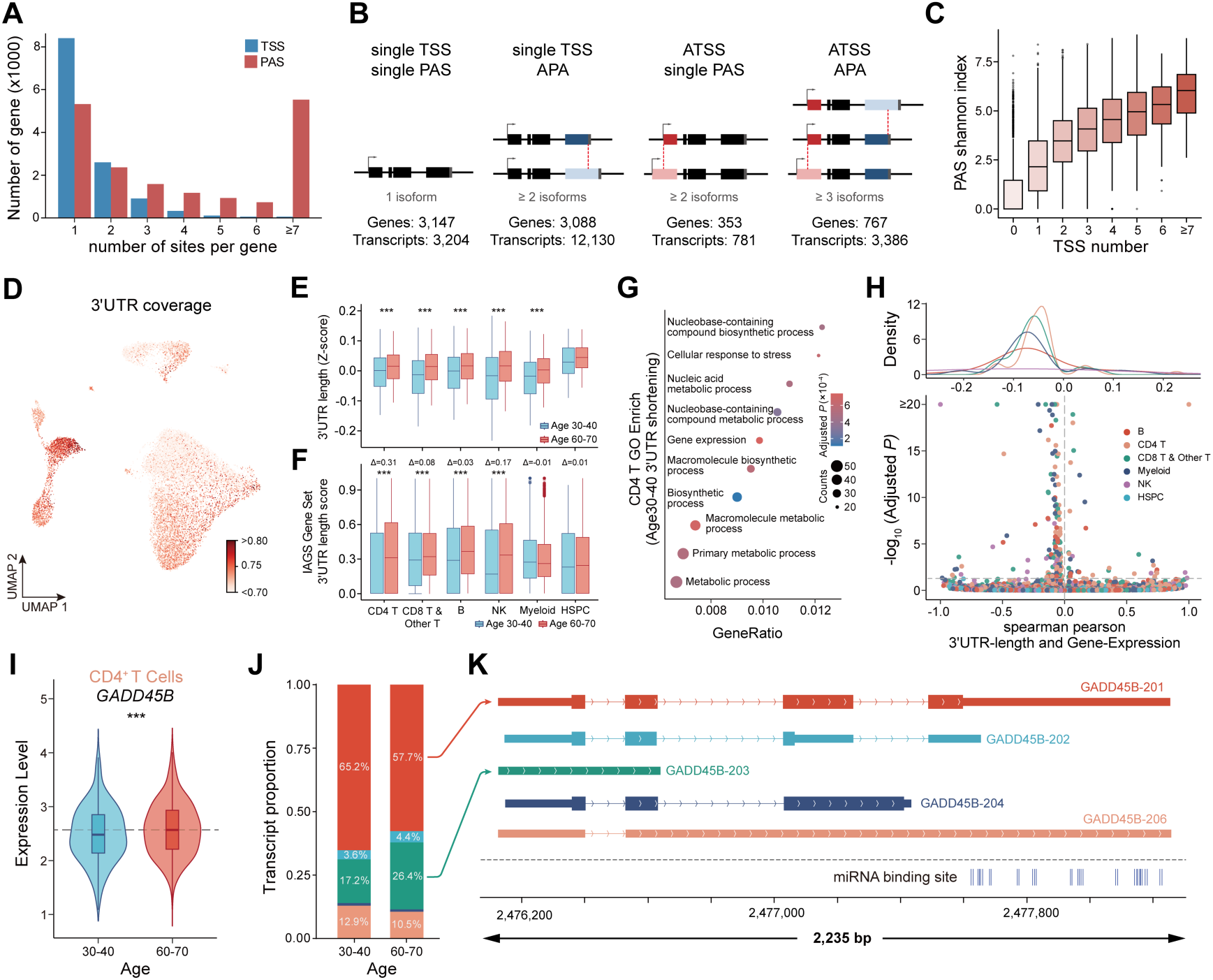
Transcript Architecture and 3′-UTR Variability Reveal Post-Transcriptional Regulation During Immune Aging. **A**, Number of transcriptions start sites (TSS, blue) and polyadenylation sites (PAS, red) per gene. **B**, Genes were categorized according to the transcription start sites (TSSs) and polyadenylation sites (PASs) detected in full-length isoforms. The presence of multiple TSSs (arrows) or PASs (stripes) marks ATSS and APA genes, respectively. **C**, 3′ end diversity, quantified using the Shannon index of all 5′-3′ isoforms, shows a positive relationship with the number of transcriptions start sites per gene, indicating that genes with more TSSs tend to exhibit greater 3′ end variability. **D**, t-SNE visualization of average 3′-UTR coverage across single cells. Each point corresponds to an individual cell, colored according to the relative 3′-UTR coverage level (white, low; red, high). **E**, Boxplots showing the distribution of 3′-UTR lengths across different immune cell subsets between age groups. Blue and red represent the 30-40-year and 60-70-year groups, respectively. Statistical significance was assessed using the Wilcoxon rank-sum test. **F**, Distribution of 3′-UTR length enrichment scores for IAGS genes across immune cell subsets in individuals aged 30-40 years (blue) and 60-70 years (red). Statistical significance was assessed using the Wilcoxon rank-sum test. **G**, Enriched GO terms associated with genes displaying significantly shortened 3′-UTRs in CD4^+^ T cells. The color scale represents enrichment significance, while the size of each dot indicates the number of genes involved. **H**, Density (top) and volcano (bottom) plots depicting the Spearman correlation between 3′-UTR length and gene expression at the single-cell resolution. **I**, Distribution of GADD45B expression in CD4^+^ T cells across the 30-40-year (blue) and 60-70-year (red) age groups. **J**, Transcript abundance of GADD45B in CD4^+^ T cells stratified by age groups. **K**, Visualization of GADD45B transcript isoforms and miRNA binding sites (miRNA annotations obtained from the TargetScan database (McGeary et al., 2019)).

As differential PAS selection directly influences 3′UTR length, we calculated the average 3′UTR coverage at the single-cell level and observed that myeloid cells exhibited the highest 3′UTR coverage among all cell types (Figure 3D; see Methods) potentially reflecting their inherently complex post-transcriptional regulatory networks as innate immune cells. We then compared UTR length between the Age 30-40 and Age 60-70 groups and observed that relative to the younger group, the older group expressed isoforms with longer 3′UTRs but shorter 5′UTRs across diverse PBMCs subsets (Figure 3E; Figure S10C-E). This age-associated lengthening of UTR can introduce additional cis-regulatory elements (Chen et al., 2018; Mitschka and Mayr, 2022), potentially activating a more intricate post-transcriptional repressive network.

To further investigate the functional implications of 3′UTR length differences, we analyzed an immunosenescence-associated gene set (IAG) (Liu et al., 2025) across different cell types. The 3′UTRs of IAGs were generally longer in the Age 60-70 group, with the most pronounced difference observed in CD4⁺ T cells (Figure 3F). We then performed functional enrichment analysis on 79 genes with significantly altered 3’UTR lengths specifically in CD4⁺ T cells (Figure S10F). These genes were highly enriched for processes related to a global decline in anabolic capacity, reduced precision of gene expression regulation, and diminished ability to cope with internal and external stress (Figure 3G). These results indicate that APA-mediated isoform selection may modulate core regulatory networks controlling cellular metabolism and homeostasis, contributing to functional deterioration in CD4⁺ T cells and providing a post-transcriptional framework for understanding immunosenescence.

Further supporting this mechanism, correlation analysis across single cells revealed a negative relationship between average 3′UTR length and gene expression, which was particularly pronounced in CD4^+^ and CD8^+^ T cells (Figure 3H). This inverse relationship was observed across numerous immune-related genes, suggesting that 3′UTR lengthening broadly affects transcript stability during aging, representative examples include GADD45B and ZFP36L2. Focusing on GADD45B, a key regulator of stress response and DNA damage repair (Liebermann and Hoffman, 2014; Rodríguez-Jiménez et al., 2021), we observed that isoforms with longer 3′UTRs contained more predicted miRNA binding sites (Figure 3I) and were associated with reduced mRNA abundance in young individuals (Figure 3J and K), implying that 3′UTR length differences may decrease mRNA stability and expression via enhanced miRNA-mediated repression (O’Brien et al., 2018). Notably, GADD45B (Moskalev et al., 2012) expression has been shown to increase upon TCR activation and to modulate effector pathways such as IFN-γ signaling (Ju et al., 2009), and its reduced expression in young individuals may therefore reflect a more tightly regulated activation threshold and genomic maintenance capacity. A similar pattern was observed for ZFP36L2, an RNA-binding protein involved in post-transcriptional regulation of immune genes, where longer 3′UTR isoforms were enriched in predicted miRNA binding sites and correlated with lower expression in aged T cells (Figure S10G-I), reinforcing the notion that 3′UTR-mediated regulation contributes broadly to immune aging. Together, these results establish a mechanistic link between transcript isoform dynamics and the functional deterioration characteristic of immunosenescence.

### Single-Cell Profiling Reveals an Aging-Driven Functional Shift in T Cells from Cytotoxicity to a Pro-Inflammatory State

We next interrogated the immune receptor repertoires to delineate how aging reshapes adaptive immune diversity. Clonal architecture reconstructed from both BCR and TCR sequencing revealed distinct aging trajectories across immune lineages. BCR repertoires maintained comparable diversity, clonality, and CDR3 length distributions across age groups and major B-cell subtypes (Song et al., 2022; Terekhova et al., 2023b). A notable, although non-significant, trend of increased clonal expansion was observed in plasma cells with age (Figure 1D and E). The only exception to this overall similarity was found in switched memory B cells, where elderly individuals exhibited slightly longer heavy chain(IGH) CDR3 regions, suggesting subtle age-related shifts in repertoire selection (Figure S11A). In contrast, T-cell repertoires demonstrated clear signs of aging-associated remodeling. Systematic analysis of 13 T cell subsets based on reconstructed TCR sequences revealed a marked contraction in repertoire diversity and increased clonality in the 60-70-year cohort, consistent with established hallmarks of immunosenescence (Luo et al., 2022) (Figure 1E). Further analysis showed that such age-related clonal expansion was not uniformly distributed across T-cell lineages but was predominantly concentrated within cytotoxic T lymphocytes (CTLs, including both CD4⁺ and CD8⁺ CTLs) cells (Britanova et al., 2016; Mogilenko et al., 2021b) (Figure 4A; Figure S11B). Quantification using STARTRAC (Zhang et al., 2018) expansion and Gini indices confirmed significantly increased clonality in aged CTLs (Figure 4B; Figure S11C), alongside a trend toward reduced diversity (Figure S11C). CDR3 length analysis demonstrated that MAIT cells exhibited a prominent peak at TCRα chian CDR3 amino acid lengths of 12-13 and preferentially utilized the TRAV1-2-TRAJ33 gene pairing, consistent with previous studies (Porcelli et al., 1993; Tilloy et al., 1999; Treiner et al., 2003), along with CTLs elongated TCRβ chain CDR3 sequences compared to younger individuals in our data (Figure 4C; Figure S11D and E). These structural features may result from chronic antigenic and inflammatory exposure, leading to reduced TCR diversity in elderly individuals and a corresponding narrowing of the antigen recognition repertoire. Collectively, these pronounced changes in the TCR landscape indicate functional remodeling of the CTL compartment during aging.

**Figure 4.**
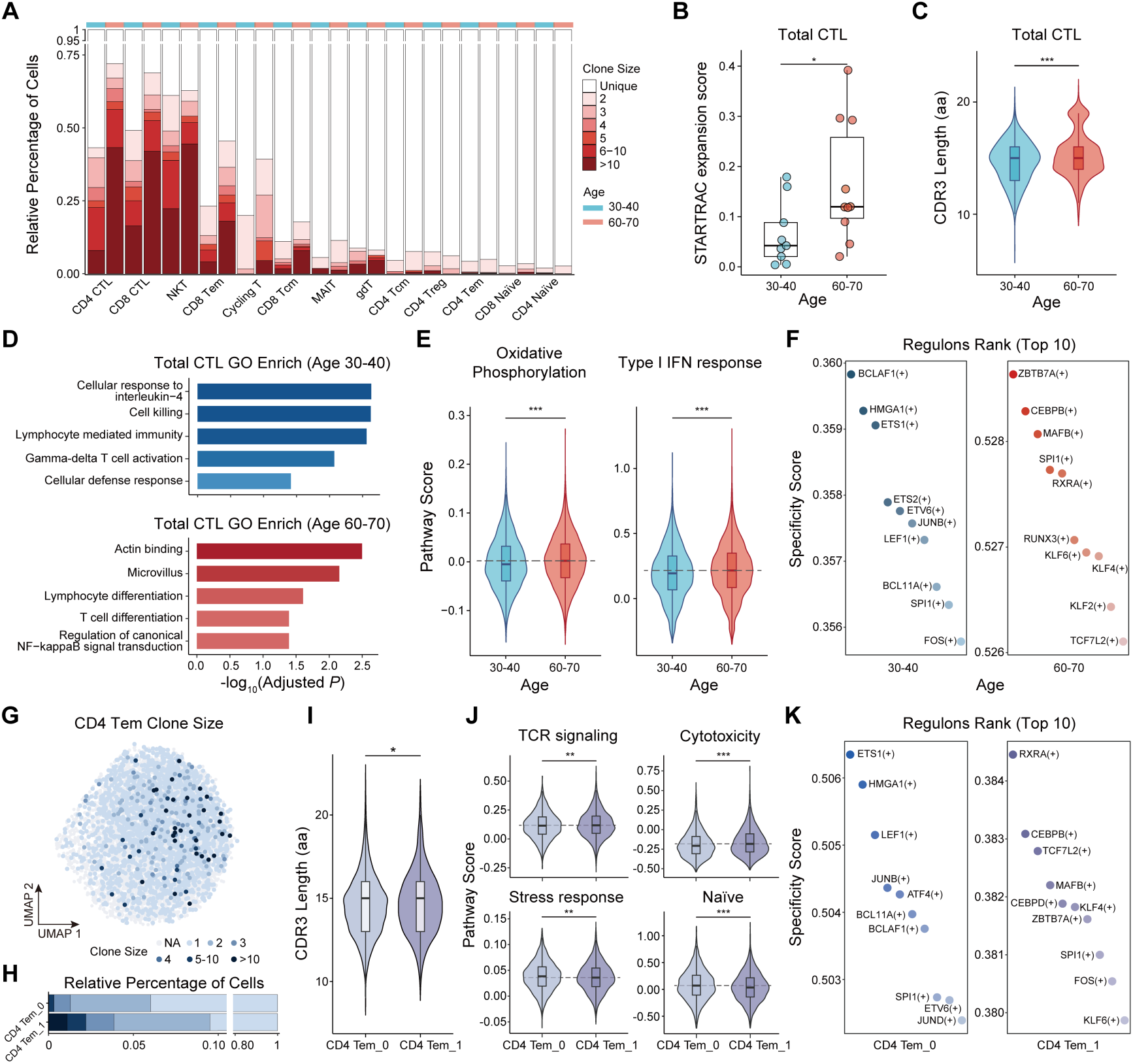
Age-related Clonal Expansion and Functional Divergence in T Cell Subsets. **A**, Clonal architecture of T-cell subsets. Effector and memory populations (e.g., CD8 CTL, CD8 Tem) show increased clonal expansion in older adults (60-70 years, light red) compared to younger (30-40 years, light blue), while Naïve populations (e.g., CD4 Naïve, CD8 Naïve) remain predominantly unique clones. **B**, STARTRAC expansion scores of total cytotoxic T lymphocytes (CTLs). Older group exhibits significantly higher clonal expansion (\**P* < 0.05, two-sided Wilcoxon rank-sum test). **C**, TRB CDR3 length distribution in total CTLs. Older adults have significantly longer CDR3 sequences (\*\*\**P* < 0.001), indicating repertoire diversification. (\**P* < 0.05, \*\**P* <0.01, \*\*\**P* <0.001, two-sided Wilcoxon rank-sum test). **D**, GO enrichment analysis for differentially expressed genes in CTL cells. Top: 30-40 age group; Bottom: 60-70 age group. Color intensity represents significance level (-log10(adjusted *P*)). **E**, Violin plots comparing pathway scores in CTL cells between younger (30-40 years) and older (60-70 years) age groups. Statistical significance of group differences in pathway scores was determined by two-sided Wilcoxon rank-sum test. **F**, Top 10 transcription factor regulons ranked by specificity score in CTL cells across age cohorts: 30-40 years (left) and 60-70 years (right). **G**, UMAP projection of CD4 Tem_0 and CD4 Tem_1 subclusters, colored by clone size. Larger clones (dark blue) localize to distinct regions of the transcriptional landscape, indicating phenotypic clustering of expanded clones. **H**, Stacked bar chart displaying the relative percentage of cells within CD4 Tem cell subsets, showing the relative frequencies of CD4 Tem_0 and CD4 Tem_1 subpopulation. The CD4 Tem_1 subset demonstrates a relatively higher proportion of clonally expanded cells. **I**, Violin plots comparing TRB CDR3 length distributions between CD4 Tem_0 and CD4 Tem_1 subsets. A significant difference (\**P* < 0.05) suggests distinct T cell receptor signatures between these functionally specialized populations. Statistical significance was determined by a two-sided Wilcoxon rank-sum test. **J**, Violin plots comparing pathway scores between CD4 Tem_0 and CD4 Tem_1 subclusters. Statistical significance of group differences in pathway scores was determined by two-sided Wilcoxon rank-sum test. **K**, Top 10 transcription factor regulons ranked by specificity score in CD4 Tem_0 subcluster (left) and CD4 Tem_1 subcluster (right).

To link TCR dynamics to transcriptional states, we analyzed the paired scRNA-seq data from the same CTLs cells, performing differential expression and functional enrichment analyses. In CTLs from the Age 30-40 cohort, upregulated genes were predominantly enriched in classical effector pathways, such as “cell killing” and “lymphocyte-mediated immunity.” In contrast, CTLs from the Age 60-70 cohort exhibited significant enrichment in inflammation-associated pathways (eg. regulation of canonical NF-κB signaling transduction) and lymphocyte differentiation pathways (Figure 4D). Additionally, the elderly cohort displayed pronounced activation of TLR signaling, type I interferon responses, cytotoxicity and oxidative phosphorylation (Figure 4E; Figure S11F). This pattern suggests a functional shift in aged CTLs from a dedicated cytotoxic program toward a state of inflammatory regulation and innate immune activation, likely driven by chronic antigenic stimulation. Analysis of transcription factor (TF) activity revealed the regulatory basis for this divergence. CTLs from the Age 60-70 cohort were characterized by high activity of ZBTB7A (a modulator of NF-κB pathways) and RUNX3 (a promoter of cytotoxic gene expression and IFN-γ production in CD8⁺ effector T cells) (Cruz-Guilloty et al., 2009; Ramos Pittol et al., 2018). By contrast, CTLs from the Age 30-40 cohort were characterized by high activity of BCLAF1, a transcriptional regulator essential for T cell activation (Kong et al., 2011) (Figure 4F).

Age-associated remodeling of clonal architecture and functional state was also observed within CD4⁺ effector memory T (Tem) cell subsets resolved by single-cell long-read sequencing. Partially due to lower expression of TRAV genes than of TRBV genes (Gao et al., 2022; Redmond et al., 2016),we analyzed the TCRβ repertoires of the CD4 Tem_0 and CD4 Tem_1 subsets. The CD4 Tem_1 subgroup, enriched in the Age 60-70 cohort (Figure 2N), showed a propensity for clonal expansion (Figure 4G; Figure S12A), reduced diversity (Figure 4H; Figure S12B), and longer CDR3 amino acid length (Figure 4I). Functionally, the CD4 Tem_1 subset was characterized by heightened TCR signaling, increased pro-apoptotic potential, and enhanced cytotoxicity, accompanied by the significant upregulation of cytokines. In contrast, the CD4 Tem_0 subset highly expressed markers of naive T cells and genes associated with stress responses, suggestive of a quiescent state with stress-adaptive potential (Figure 4J; Figure S12C) (Hamilton and Jameson, 2012). Regulatory network analysis further supported these distinctions: RXRA activity was most prominent in CD4Tem_1, whereas ETS1 dominated in CD4Tem_0 (Figure 4K), revealing at the transcriptional regulatory level that CD4Tem_1 is skewed toward an effector differentiation state, while CD4Tem_0 tends to maintain quiescence and homeostasis (Garrett Sinha, 2023; Stephensen et al., 2007; Zhou et al., 2023). Collectively, these results reveal subset-specific transcriptional and clonal patterns in aging, supporting distinct functional properties across these subsets.

## DISCUSSION

Aging is not only characterized by the gradual decline of organ function but also, more fundamentally, by progressive impairment of the immune system, a phenomenon known as immunosenescence. To dissect the molecular and cellular underpinnings of this phenomenon, we employed an integrative multi-omics approach, combining short-read and long-read single-cell transcriptomics with immune receptor profiling. This strategy enabled the construction of a comprehensive atlas capturing gene expression patterns, transcript isoform diversity, and immune receptor repertoires at single-cell resolution. By leveraging this dataset, we moved beyond conventional analyses of gene expression abundance to directly link cellular clonal identity, transcript isoform diversity, and functional states. This approach allowed us to uncover the complex regulatory layers underlying immunosenescence.

Our analysis systematically delineates and validates the multilayered features of immunosenescence at single-cell resolution. Beyond confirming classical observations, such as the depletion of naïve CD8⁺ T cells and expansion of regulatory T cells in elderly individuals, we uncovered extensive differences in transcript isoform abundance and usage that extends beyond changes in gene expression. Our analysis systematically delineates and validates the multilayered features of immunosenescence at single-cell resolution. Beyond confirming classical observations, such as the depletion of naïve CD8⁺ T cells and expansion of regulatory T cells in elderly individuals, we uncovered extensive differences in transcript isoform abundance and usage that extends beyond changes in gene expression. Importantly, our data reveal a coherent mechanistic model in which clonal selection in T cells and differential isoform usage act in concert to drive functional deterioration during aging.

Importantly, our data reveal a coherent mechanistic model in which clonal selection in T cells and differential isoform usage act in concert to drive functional deterioration during aging. TCR repertoire analysis demonstrated a pronounced reduction in immune diversity in the elderly, accompanied by strong clonal expansion of specific cytotoxic T lymphocyte (CTL) clones. Critically, these expanded clones did not retain classical effector functionality; instead, their transcriptional programs shifted systematically away from cytotoxic pathways toward an inflammatory state characterized by NF-κB and type I interferon signaling. These findings provide compelling evidence that chronic antigen stimulation during aging not only reduces immune diversity quantitatively but also qualitatively reprograms key effector cells toward a pro-inflammatory state.

We further identified two key transcript-level regulatory mechanisms driving this functional reprogramming. The first involves cell type-specific isoform usage. In CD4⁺ effector memory T cells, we identified a subpopulation enriched in elderly individuals (CD4 Tem_1) characterized by increased expression of truncated HSPA5 isoforms. Considering HSPA5’s function in T-cell activation, the expression of these specific isoforms may weaken cellular stress responses, increasing susceptibility to senescence, which is consistent with the clonal expansion observed in this subpopulation. The second mechanism represents a broader post-transcriptional regulatory layer, manifested as differential 3′UTR lengths. Notably, in CD4⁺ T cells, which play a central role in aging, differential 3′UTR lengths were associated with altered expression of immune-related genes. This association may be mediated by miRNA binding, as longer 3′UTRs increase the number of potential miRNA target sites, potentially reducing mRNA stability and translational efficiency. Collectively, from specific isoform usage differences to systematic 3′UTR variation, these transcript-level changes provide a mechanistic framework underlying the functional decline and phenotypic skewing of T cells during immunosenescence.

Despite providing valuable insights, this study has several limitations. First, our analyses were limited to peripheral blood cells; future studies should extend to tissue-resident immune populations to investigate aging features within specific microenvironments. Second, given that our study provides a single-timepoint snapshot of individuals, it cannot capture the dynamic trajectory of aging. Longitudinal multi-omics profiling of the same individuals over extended periods would enable a more precise characterization of immunosenescence. Finally, the functional consequences of the transcript-level alterations identified here, such as HSPA5 isoform changes and 3′UTR variation, remain to be directly validated through functional experiments, including CRISPR-based perturbations.

In conclusion, by constructing an unprecedented, multi-dimensional atlas of the aging immune system, our study systematically elucidates the pivotal role of transcript isoforms and post-transcriptional regulation in shaping the immunosenescent phenotype. These findings not only enhance mechanistic insights into aging at the molecular and cellular levels but also establish a valuable multi-dimensional resource for the research community, facilitating future studies on immunosenescence.

## MATERIALS AND METHODS

### Participants and Ethics Statement

This study utilized data from the I99 Health Cohort, a prospective multi-omics population study initiated in 2024. Participants aged 18-99 years were recruited from health examination centers in Shenzhen, Wuhan, Hangzhou, and Qingdao, with planned 5-year follow-ups including longitudinal biosample collection and multidimensional health data acquisition. The cohort aims to establish dynamic health baselines across life stages and investigate age-related variations in multi-omics profiles.

For the present analysis, we selected two age groups: (1) Age30-40 (n=9, 4 males/5 females) and (2) Age60-70 (n=10, 5 males/5 females). All participants were free of major diseases (e.g., malignancies, severe cardiovascular events, or stroke sequelae) and had no history of transplants, stem cell therapy, allogeneic transfusion, or exogenous DNA-based immunotherapies.

The study protocol was approved by the Institutional Review Board of BGI (Approval No: BGI-IRB 23123-T1). All participants provided written informed consent, and the study adhered to all relevant human research ethics regulations.

### Collection of Peripheral Blood Samples and Cell Cryopreservation/Revival

Peripheral blood (10 mL) was collected from the antecubital vein of healthy volunteers into EDTA-coated tubes. Peripheral blood mononuclear cells (PBMCs) were isolated using Ficoll density gradient centrifugation. Briefly, whole blood was centrifuged at 1,600 ×g for 10 min, after which the plasma was removed and replaced with an equal volume of PBS. The diluted blood was carefully layered onto Ficoll solution and centrifuged at 800 ×g for 20 min. The buffy coat was collected, washed twice with PBS, and resuspended. Cell viability was assessed by trypan blue staining. The cell pellet was centrifuged at 500 ×g for 5 min and resuspended in cell freezing medium (FBS:DMSO = 9:1, v/v). The cells were cryopreserved via programmed freezing and stored in liquid nitrogen.

For revival, cryovials were rapidly thawed in a 37°C water bath. The cell suspension was transferred into RPMI-1640 medium supplemented with 10% FBS, mixed by gentle inversion, and centrifuged at 300 ×g for 10 min. The supernatant was discarded, and the cells were resuspended in PBS containing 0.04% BSA. Cell viability and count were re-evaluated before subsequent experiments.

### Single-cell RNA Library and TCR Library Preparation and Sequencing

Single-cell RNA sequencing of the PBMC suspension was performed using the DNBelab C4 system. Specifically, the DNBelab C series high-throughput single-cell 5′ RNA and V(D)J library preparation set was used in combination with a pressure-driven single-cell droplet generator. The sample suspension, barcoded beads, and oil phase were mixed on an scRNA chip to generate single-cell emulsion droplets. Within the droplets, cell lysis was performed, and the released mRNA was labeled with barcodes on the beads. This was followed by emulsion breaking, filtration, and reverse transcription to synthesize full-length cDNA. The resulting cDNA was divided into two parts: one half was used to construct short-read scRNA-seq and TCR-seq libraries, which were sequenced on the BGISEQ-T1 platform; the other half was used to prepare long-read scRNA-seq libraries, which were sequenced on the CycloneSEQ G400-ER sequencer.

### Single-cell short-read data preprocessing

The reference genome sequence (GRCh38.primary_assembly.genome.fa) and genome annotation file (gencode.v32.primary_assembly.annotation.gtf) for Homo sapiensused in this study were downloaded from the GENCODE database (Release 32). Subsequently, the reference genome index was constructed using the BGI STOmics Cloud pipeline scRNA-seq_C4_analysis_pipeline_build_index (v3.1.5). The raw data were processed using the scVDJ-seq pipeline (v1.1.0) to obtain the single-cell gene expression matrix and barcode information.

The single-cell expression matrix was constructed and quality control analysis performed using the Seurat R package (v4.3.0, based on R v4.2.2 environment)(Hao et al., 2021; Satija et al., 2015). Cell filtering criteria were as follows: the number of genes detected per cell had to be between 400 and 5,000, the mitochondrial gene percentage had to be below 10%, and the hemoglobin gene percentage had to be less than 1.5%. Subsequently, the DoubleFinder package was used to estimate doublet rates (calculated as total cells × 8 × 10⁻⁶) and remove doublets (McGinnis et al., 2019). To further enhance data quality, the following genes were excluded from subsequent analyses: ribosomal genes, mitochondrial genes, pseudogenes and long intergenic non-coding RNAs (lincRNAs). Genes encoding α and β TCR chains, heavy and light chains of BCR, and histone genes were excluded from the subsequent dimensionality reduction clustering and cell annotation analysis.

### Dimension Reduction and Batch Effect Correction

Based on quality-controlled data, we performed single-cell transcriptomics analysis using the Seurat package (Hao et al., 2024). First, gene expression was normalized using the LogNormalize method, and highly variable genes were identified with default parameters of FindVariableFeatures. Subsequently, the data were scaled and subjected to linear dimensionality reduction via principal component analysis (PCA) using the ScaleData and RunPCA functions. To eliminate batch effects, the Harmony package was utilized to perform integrated correction on the principal components (Korsunsky et al., 2019). Finally, based on the corrected results, the FindNeighbors and FindClusters functions were used for cell clustering, followed by nonlinear dimensionality reduction and visualization via UMAP.

### Cell Annotation

We first performed unsupervised clustering of all cells and identified seven major cell types based on classical marker genes and inter-cluster differentially expressed genes (identified using the FindMarkers function): CD4⁺ T cells, CD8⁺&other T cells, natural killer (NK) cells, B cells, hematopoietic stem and progenitor cells (HSPCs)&myeloid cells, and platelets. Platelets were excluded from subsequent analyses.

To further resolve cellular heterogeneity, we performed subcluster-level re-clustering for CD4⁺ T cells, CD8⁺ and other T cells, NK cells, B cells, and HSPCs&myeloid cells. During this process, cells expressing signature genes from two or more major cell types were defined as “twins” and removed from analysis. Ultimately, we defined 30 cell subpopulations: 5 CD4⁺ T cell subsets, 8 CD8⁺ and other T cell subsets, 4 NK cell subsets, including 5 B cell subsets, 8 HSPCs&myeloid cell subsets, and 1 HSPC subset. Characteristic gene lists for each subpopulation are detailed in the Supplementary Tables.

### Differentially Expressed Gene Analysis and GO Enrichment Analysis

To identify differentially expressed genes (DEGs) between the two groups, we employed the “FindMarkers” function in the Seurat R package. Genes were marked as significant DEGs if they met the following criteria: adjusted *P* < 0.05, aan absolute average log₂(fold change) > 0.15, expression in more than 10% of cells in either group (pct > 0.1). Gene Ontology (GO) enrichment analysis of the DEGs was performed using the clusterProfiler (Wu et al., 2021) package (v4.14.6) to compare functional enrichment in biological processes (BP) across different age groups (Wu et al., 2021). *P* were corrected using the Benjamini-Hochberg (BH) method, with significance thresholds set at *P* < 0.05 and q < 0.05.

### Module Score Assessment for Cells

We assessed the module scores for each cell using the “AddModuleScore” function from the Seurat R package. The gene sets utilized in this analysis were obtained from previously published studies, including the immunosenescent-associated gene (IAG) set, ImmuneSigDB (Godec et al., 2016), and gene sets related to CD4^+^ T cell states (Chu et al., 2023).

### Single-Cell Regulatory Network Analysis and Transcription Factor Activity Quantification

To decipher gene regulatory networks in PBMCs, we employed the SCENIC workflow (Aibar et al., 2017). First, we constructed gene co-expression networks using the GRNBoost2 algorithm. Subsequently, motif enrichment analysis was performed using the cis-regulatory element database (hg38) to identify candidate regulators, and AUCell was applied to calculate regulator activity at the single-cell level. Finally, we screened transcription factors specifically activated in each cell type or group using regulator specificity scores (RSS), thereby systematically revealing cell type-specific or group-specific transcriptional regulatory programs.

### Single-cell long-read data preprocessing

Raw FASTQ reads were first processed with Seqkit (Shen et al., 2016) to remove fragments shorter than 300 bp. Next, Fastp (Chen, 2023) was used to perform quality control and filter out low-quality reads. Adapter-containing reads were then retained using Cutadapt (Marcel, 2011), excluding reads without the expected adapters. Barcode demultiplexing was carried out by aligning reads to a reference barcode list, and reads matching a valid barcode were retained(Lin et al., 2024). These reads were then mapped to the human genome (GRCh38) using Minimap2 (Li, 2018) in a splice-aware mode to generate alignment files. UMI correction was applied using UMI-tools (Smith et al., 2017). IsoQuant (Smith et al., 2017) generated files that were used in downstream analyses. Finally, transcripts were classified and annotated using SQANTI3 (Pardo-Palacios et al., 2024).

### Analyses of Single-cell Long-read Transcript Expression

We began with the output from IsoQuant, using raw transcript quantification for single-cell long-read data. To ensure robustness of the dataset, we applied stringent filtering criteria: cells with fewer than five detected transcripts were excluded; transcripts detected in fewer than three cells were removed; and transcripts absent from at least three different individuals were also discarded. Cell cluster labels from short-read data were assigned to corresponding long-read cells via cell IDs. To mitigate sparsity in the resulting expression profiles, we applied SEACells [73] to aggregate cells into metacells. Within each cell type, cells from the same individual were grouped into metacells according to expression similarity, with an approximate aggregation ratio of 50 cells per metacell.

The final metacell-level transcript expression matrix was log-normalized, and dimensionality reduction was performed using Seurat’s CCA integration to align variation across individuals and cell types. Separately, cell state identification was conducted using the original transcript expression matrix within major cell groups.

### Differential transcript usage (DTU) analysis

To evaluate DTU between consecutive developmental stages, we employed IsoformSwitchAnalyzeR (Vitting-Seerup and Sandelin, 2019). Transcript expression values were aggregated into bulk-level matrices by concatenating sample and fine-grained cell type identifiers (e.g., sample_L3celltype). DTU comparisons were then performed between groups, applying a statistical cutoff of *P* < 0.05 and an absolute difference (|dIF|) greater than 0.1.

### Identification of TSS and PAS and Quantification of UTR Lengths at Single-Cell Resolution

We first identified the start and end positions of each read in the sequencing data. To minimize artifacts introduced during reverse transcription in A-rich regions, reads ending within such regions were filtered out to avoid spurious signals. For transcription start site (TSS) identification, we validated candidate sites against the refTSS database (Abugessaisa et al., 2019) and retained only those located within ±50 bp of annotated TSSs.

After obtaining high-confidence TSSs and polyadenylation sites (PASs), we compared them with the start and stop codons of the corresponding genes to infer the UTR length and coverage of each transcript. Based on this, we calculated the expression-weighted UTR length (EUL) for each gene in each cell. Specifically, UTR lengths of all identified transcripts of a given gene were averaged after weighting by their expression levels, ensuring that highly expressed isoforms contributed more to the final EUL and providing a more accurate representation of the UTR landscape in each cell.

At the gene level, we normalized UTR lengths across reads of the same gene using z-score scaling and applied a linear mixed-effects model (LMM) to identify genes with differential UTR usage. Experimental batch was included as a random effect to account for potential batch-specific biases. Genes were defined as exhibiting significant UTR shortening or lengthening when the coefficient estimate was less than 0 (shortening) or greater than 0 (lengthening), with a corresponding P value < 0.05 and a UTR z-score difference greater than 0.1 (lengthening) or less than -0.1 (shortening).

### Single-cell TCR and BCR receptor profiling

T cell receptor (TCR) sequencing for each sample was performed using the filtered_contig_annotations.csv files generated by BGI’s DCS Cloud (scVDJ-seq v1.2.0). Data processing and analysis were conducted with the scirpy (Sturm et al., 2020), scRepertoire (Borcherding et al., 2020) and dandelion (Suo et al., 2024) toolkits. To ensure reliable clonotype identification, our analysis included only T cells possessing at least one productive TCR α chain and one productive TCR β chain. For T cells with multiple assembled α or β chains, the chain with the highest expression (based on UMI count or read depth) was designated as the dominant chain for subsequent analysis. A TCR clonotype was defined by a unique paired amino acid sequence of the TCR α and TCR β chains within a single cell.

For B cell receptor (BCR) profiling, clonotypes were identified based on a unique paired heavy chain (IGH) and light chain (IGL or IGK) combination, considering both the CDR3 nucleotide sequence and the rearranged VJ genes. A threshold for clonal grouping was calculated using the hh_s5f model distance via ddl.pp.calculate_threshold(). Only B cells with at least one productive IGH chain and one productive light chain were retained for further analysis.

The identified TCR and BCR sequences, along with their corresponding cell barcodes, were then integrated with the cell barcodes from the single-cell RNA sequencing library of the same sample. Downstream analysis was restricted to cells with productive immune receptor sequences. Clonal calling for TCRs was performed using the amino acid sequence of the CDR3 region (cloneCall=“aa” in scRepertoire functions), while BCR clonal calls utilized a “strict” definition (cloneCall=“strict”), which incorporates both the nucleotide sequence and V(D)J gene usage.

### Repertoire diversity analysis

To assess the clonal expansion of TCR and BCR repertoires, we utilized the scRepertoire package. Clonal diversity was quantified using the Shannon, Inv.simpson and Chao1 index, measures of entropy calculated with the formula: The Shannon index (H) measures repertoire entropy, integrating both clonal richness and evenness:

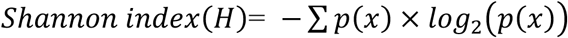

The Inverse Simpson index emphasizes clonal dominance, reflecting the extent to which the repertoire is skewed toward expanded clones:

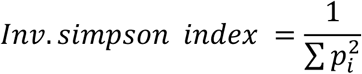

The Chao1 index estimates the total clonotype richness by accounting for unobserved, low-frequency clones:

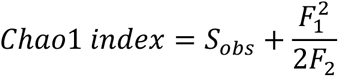

where *p(x)* represents the relative frequency of a given productive TCR or BCR clone among all αβ T cells or B cells, *Sobs* is the number of observed clonotypes, *F1* and *F2* denote the counts of singleton and doubleton clones.Together, these indices provide a comprehensive and quantitative framework for assessing immune repertoire diversity and clonal expansion, enabling robust comparisons across different cell subsets, age groups, or disease states.

### Statistics and Reproducibility

To clearly present research data, we employed multiple visualization and statistical methods. Gene expression patterns were depicted using scatter plots and heatmaps; cell proportions were displayed via box plots; and the distributions of pathway activity and gene expression were shown using violin plots. For group comparisons, the Wilcoxon rank-sum test was applied, and the resulting P-values were adjusted for multiple testing using the Bonferroni correction. Only statistically significant results are annotated in figures: *P* ≤ 0.05, *P* ≤ 0.01, and *P* ≤ 0.001 are marked with *, **, and ***, respectively. Results with *P* > 0.05 or adjusted *P* > 0.05 were considered non-significant and were not annotated.

## Acknowledgements

We sincerely thank all participants for their invaluable contributions and our team members for their support.

## Author Contributions

C.L., Y.Z., Z.R. and J.Y. conceived the idea; C.L. supervised the work; C.L., Y.Z. and J.Y. designed the experiment; Y.Y., Y.H. and W.Z. performed the majority of the experiments; Y.Z., Z.R., Y.Y., W.L. and F.L. analyzed the data; Y.Z., T.Z., X.W., X.L. and W.L. provided technical support; X.Z., B.G., Y.F., X.Y., X.L., X.J., H.L. Y.D., X.Y. and X.X. gave the relevant advice; Z.R., Y.Z., Y.Y., W.L., F.L. and C.L. wrote the manuscript.

---

**Figure S1.**
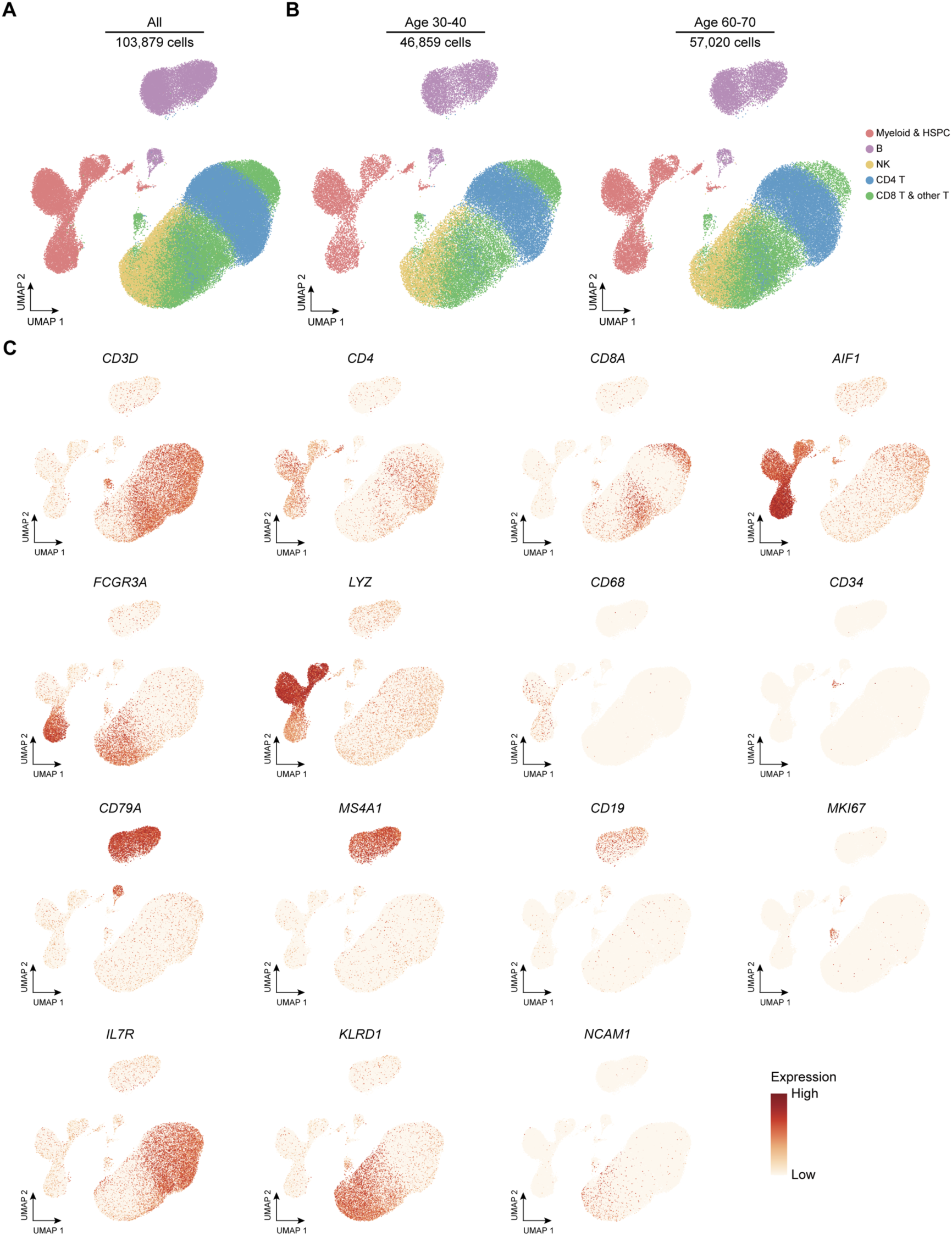
Clusters of Major Immune Cell Populations in the Age 30-40 Group and the Age 60-70 Group Derived from scRNA-seq Data. **A**, UMAP visualization of all single cells based on the scRNA-seq data. **B**, UMAP plot showing major cell types for the two sample groups. **C**, UMAP projection of cells, with expression levels of canonical markers (e.g., *CD3D*, *CD4*, *CD8A*, *AIF1*, *FCGR3A*, *LYZ*, *CD14*, *CD34*, *CD79A*, *MS4A1*, *CD19*, *MKI67*, *IL7R*, *KLRD1*, *NCAM1*) shown in color.

**Figure S2.**
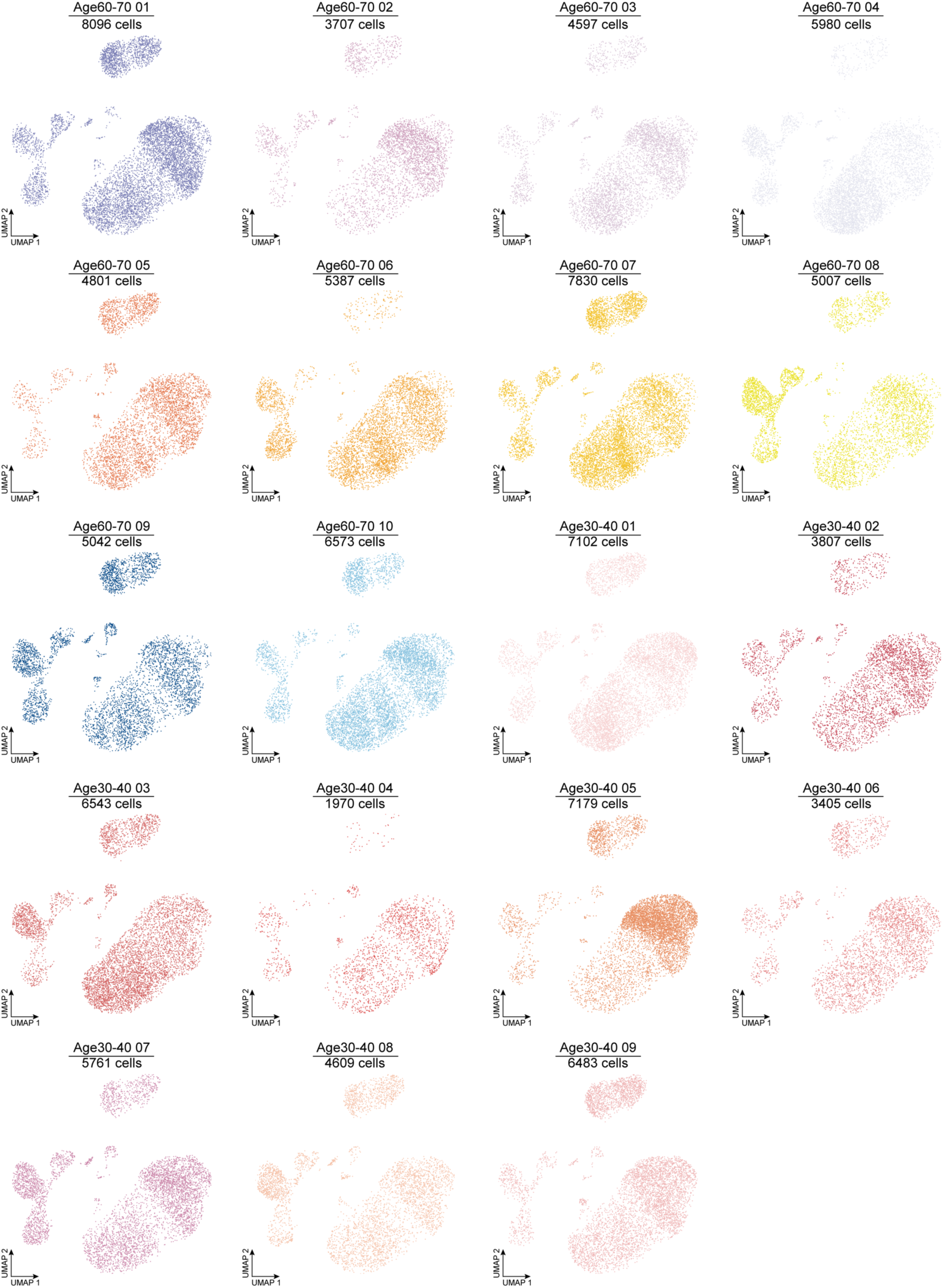
Separate UMAP Plots for Each Donor.

**Figure S3.**
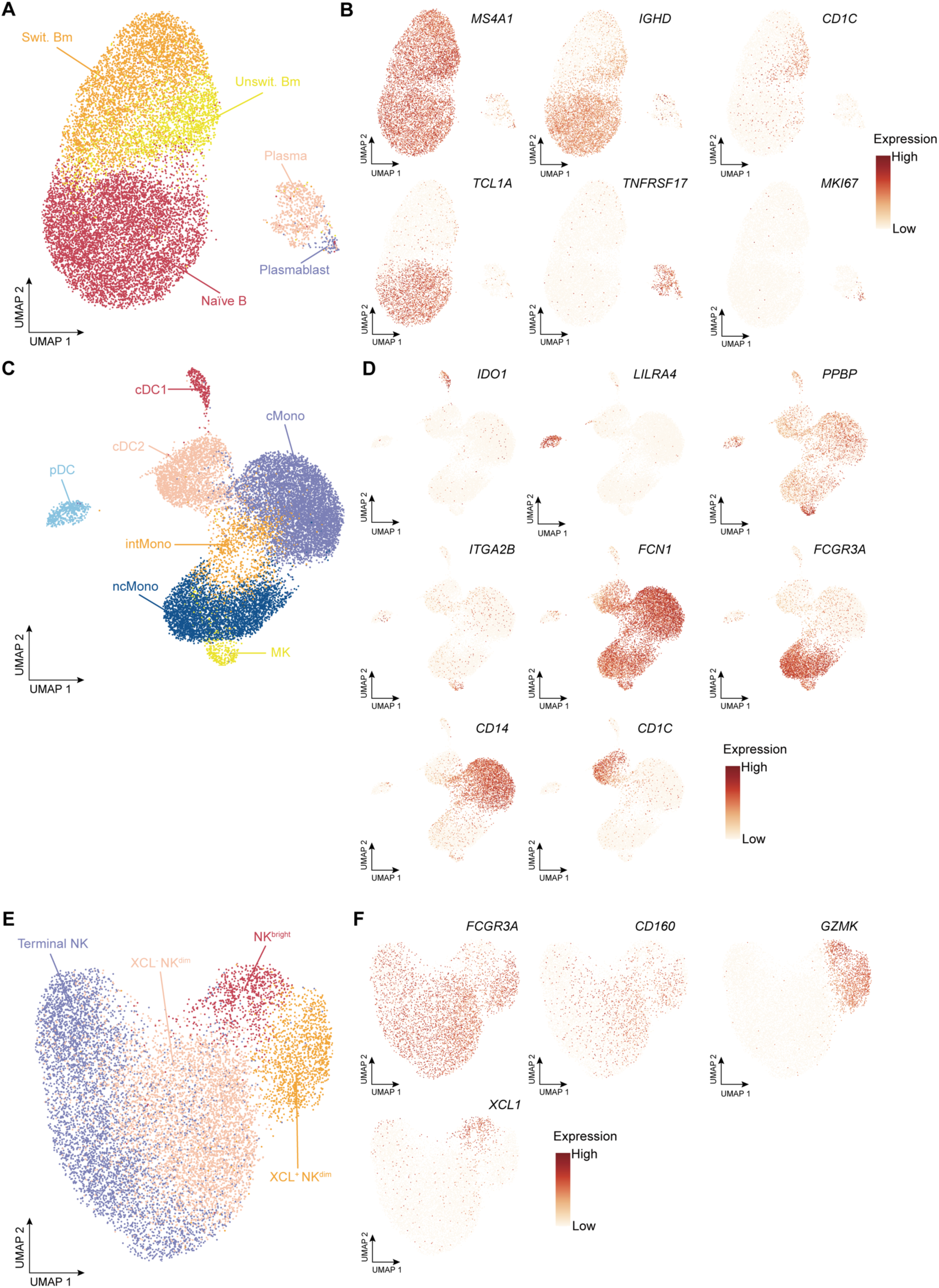
Subclustering of B and Plasma Cell, HSPC and Myeloid Cell and NK Cell Subsets Derived from scRNA-seq Data. **A**, UMAP projection of B and Plasma cell subsets. **B**, Feature plots of canonical markers for B and Plasma cell subsets. **C**, UMAP projection of Myeloid cell subsets. **D**, Feature plots of canonical markers for Myeloid cell subsets. **E**, UMAP projection of NK cell subsets. **F**, Feature plots of canonical markers for NK cell subsets.

**Figure S4.**
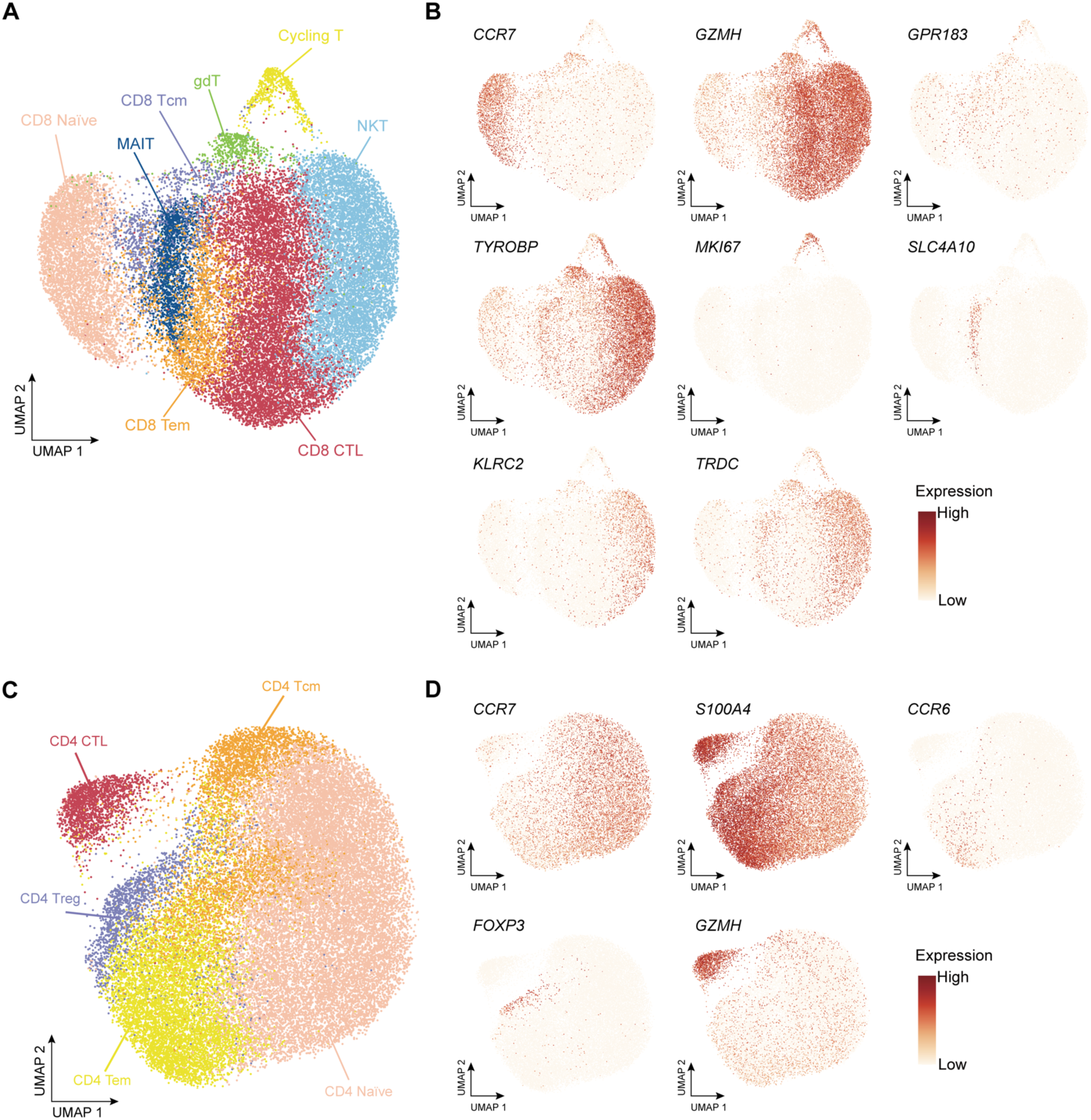
Subclustering of CD8 T & other T cell and CD4 T cell subsets derived from scRNA-seq data. **A**, UMAP projection of CD8 T & other T cell subsets. **B**, Feature plots of canonical markers of CD8 T & other T cell subsets. **C**, UMAP projection of CD4 T cell subsets. **D**, Feature plots of canonical markers of CD4 T cell subsets.

**Figure S5.**
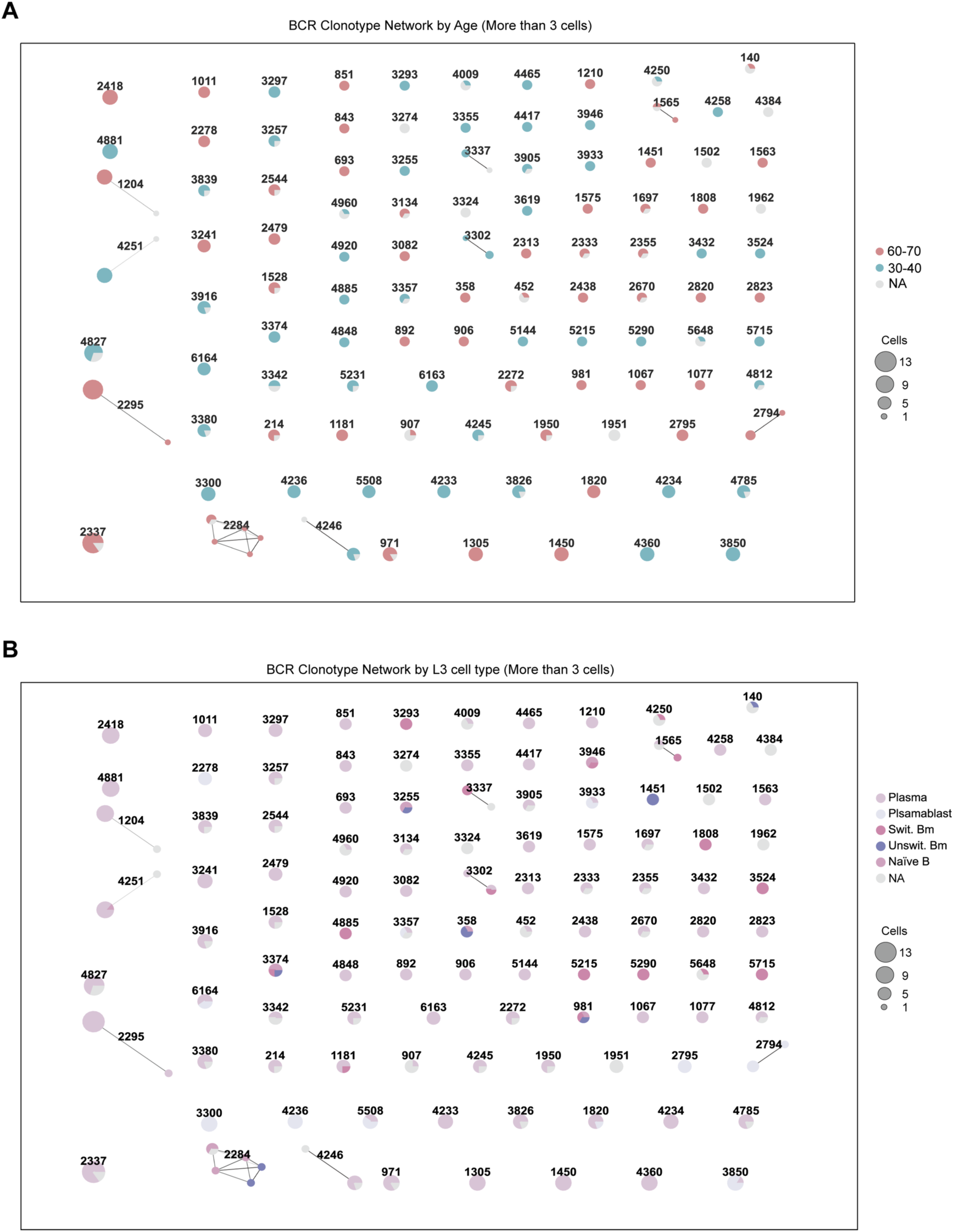
BCR Clonotype Network. **A**, Network analysis of BCR clonotypes reveals age-associated restructuring of the B cell repertoire between individuals aged 30-40 and 60-70 years. **B**, Network analysis of BCR clonotypes stratified by L3 cell type reveals connectivity patterns among distinct B-cell subsets.

**Figure S6.**
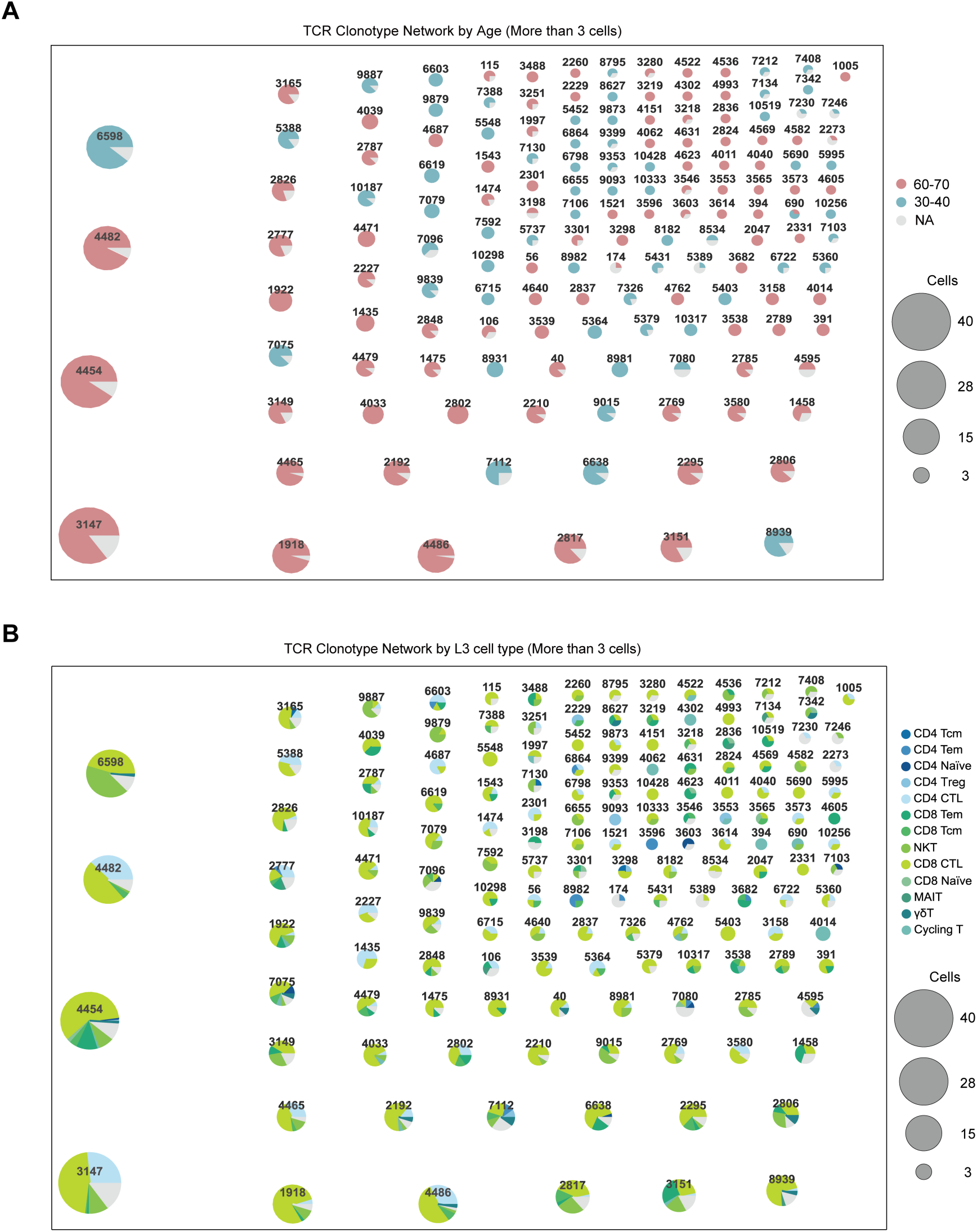
TCR Clonotype Network. **A**, Network analysis of TCR clonotypes reveals age-associated restructuring of the T cell repertoire between individuals aged 30-40 and 60-70 years. **B**, Network analysis of TCR clonotypes stratified by L3 cell type reveals connectivity patterns among distinct T-cell subsets.

**Figure S7.**
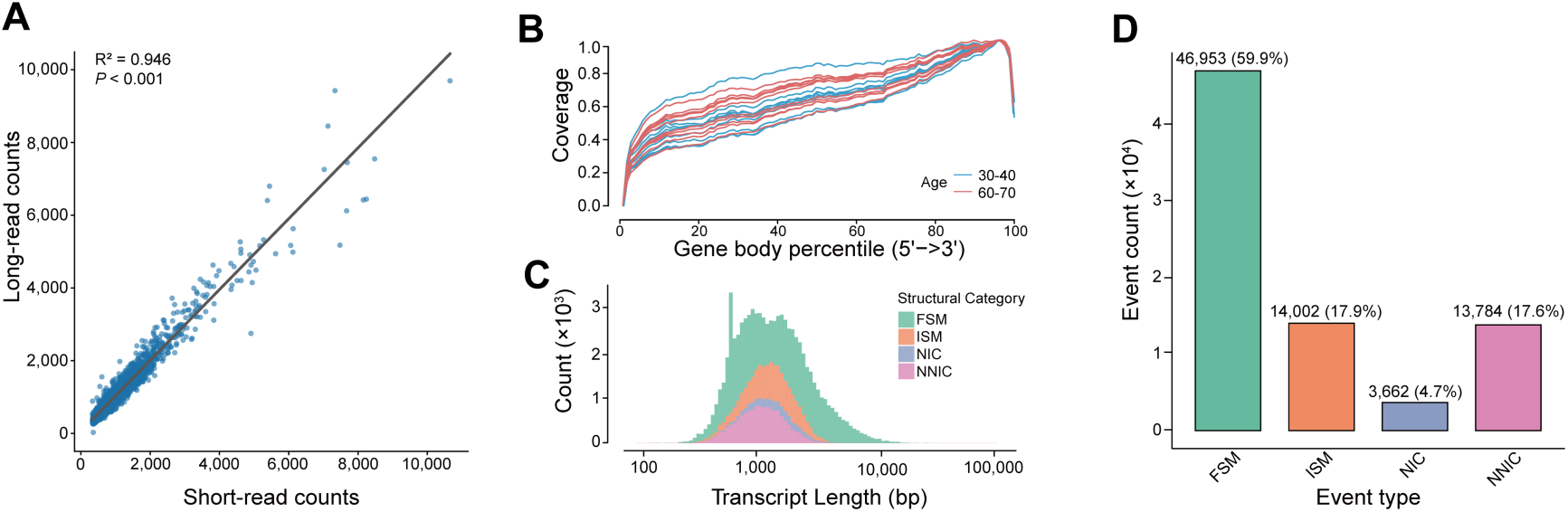
Overview and Quality Assessment of Full-Length Transcript Profiling. **A**, Scatter plot comparing per-cell UMI counts from long-read (y-axis) and short-read (x-axis) sequencing for a representative sample (Age60-70_2). Each point represents a single cell, demonstrating the high correlation in gene expression quantification between the two methods (R² = 0.946). **B**, Gene body coverage distribution across age groups (blue: 30-40 years; red: 60-70 years). **C**, Stacked histogram showing the transcript length distribution for each isoform category. The x-axis represents the transcript length in base pairs (bp) on a log scale, and the y-axis indicates the transcript count. **D**, Bar chart illustrating the distribution of identified transcript isoforms based on their comparison to existing annotations. Categories include Full Splice Match (FSM), Incomplete Splice Match (ISM), Novel In Catalog (NIC), and Novel Not in Catalog (NNC).

**Figure S8.**
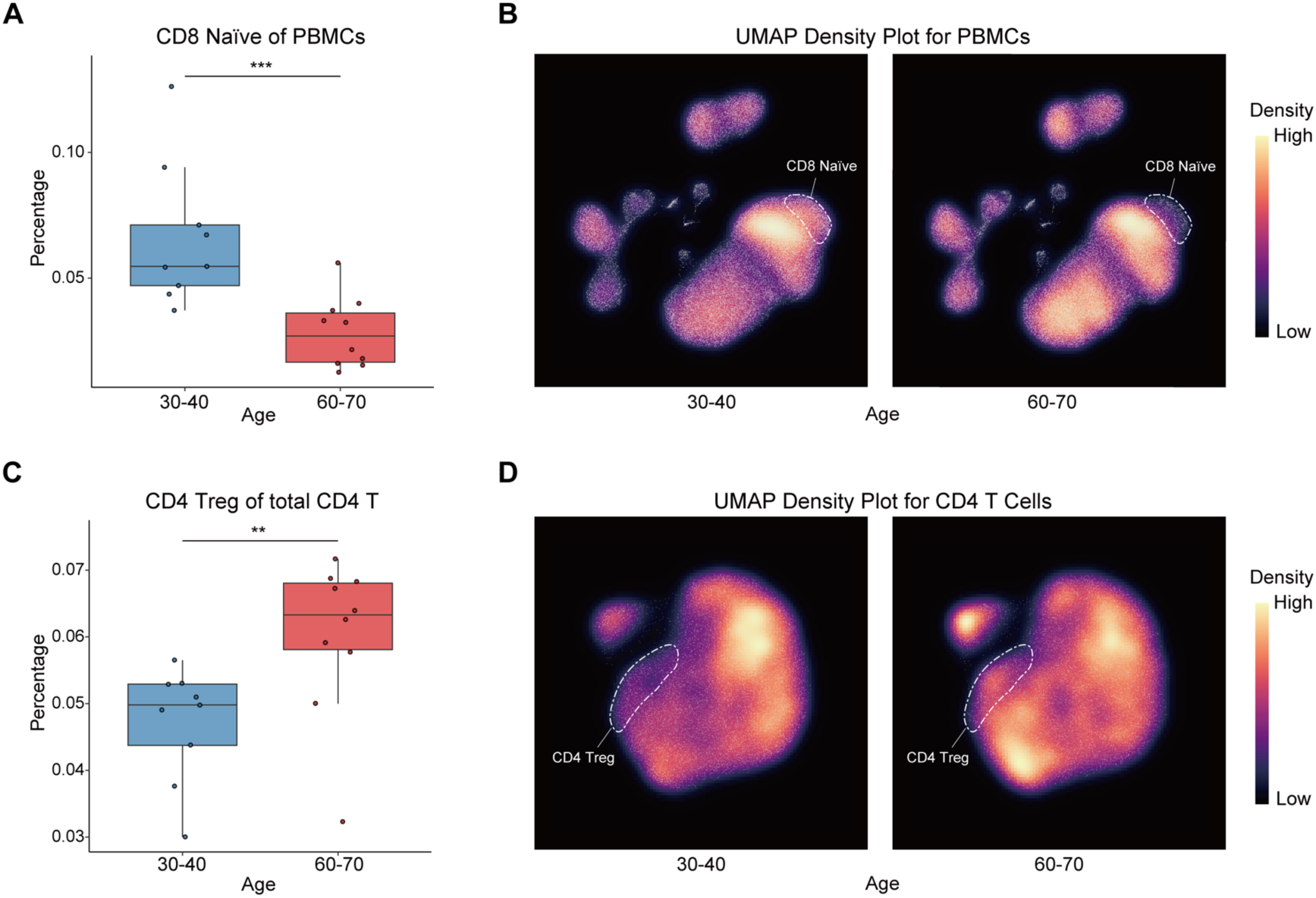
Changes in Cell Proportions Associated with Aging. **A**, Comparison of the percentage of CD8 naïve T cells in PBMCs between the Age 30-40 group (n=9) and the Age 60-70 group (n=10). **B**, Density distribution of PBMCs from the Age 30-40 group and the Age 60-70 group. **C**, Comparison of the percentage of CD4 regulatory T (Treg) cells in PBMCs between the Age 30-40 group (n=9) and the Age 60-70 group (n=10). **D**, Density distribution of CD4^+^ T cells from the Age 30-40 group and the Age 60-70 group.

**Figure S9.**
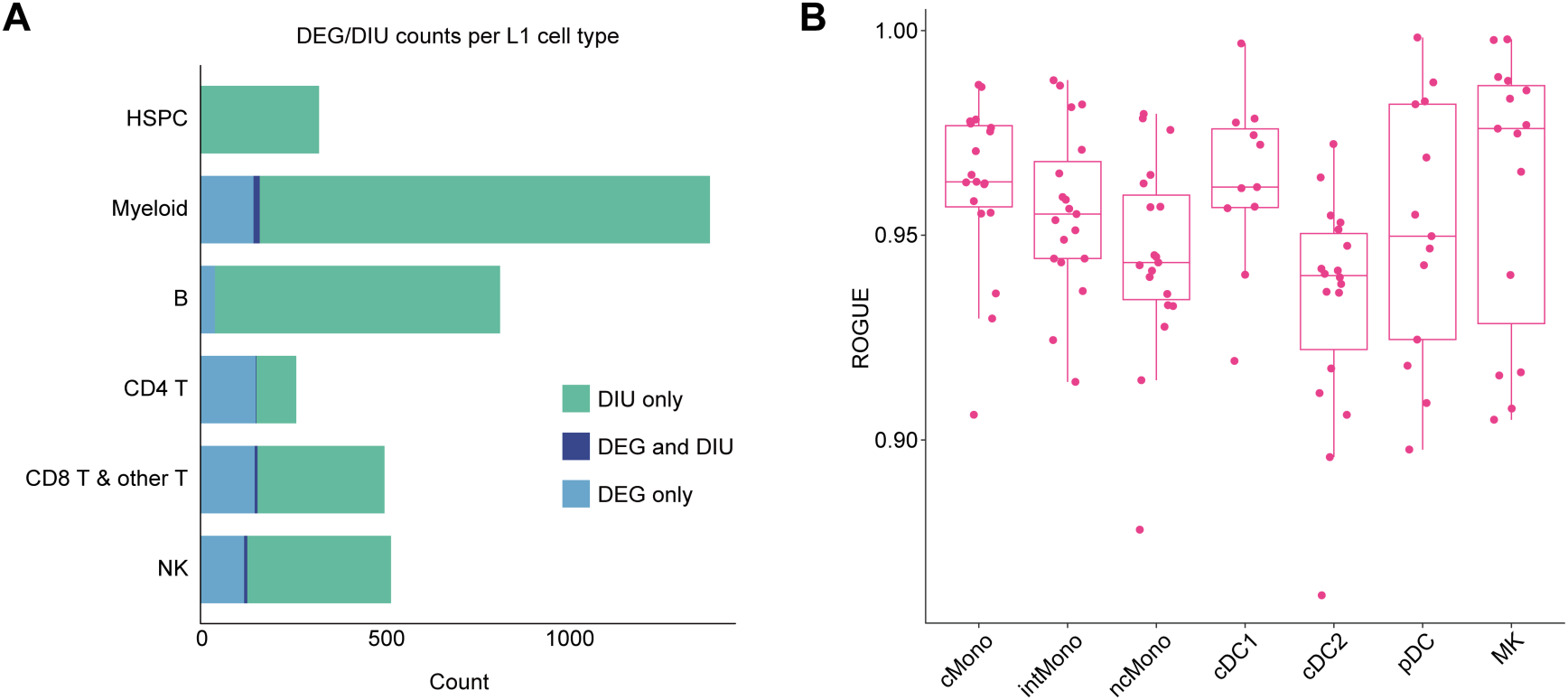
Quantification of Differential Gene and Isoform Usage and Assessment of Myeloid Subset Heterogeneity. **A**, Stacked bar chart showing the number of genes identified as differentially expressed (DEG), having differential isoform usage (DIU), or both, across major (L1) immune cell types. **B**, Box plot displaying the ROGUE (Ratio of Global Unshifted Entropy) scores for myeloid subsets. The ROGUE score is used to assess intra-cluster purity, where a higher score indicates a more homogeneous cell population (i.e., lower heterogeneity). Each point represents an individual sample within the cluster.

**Figure S10.**
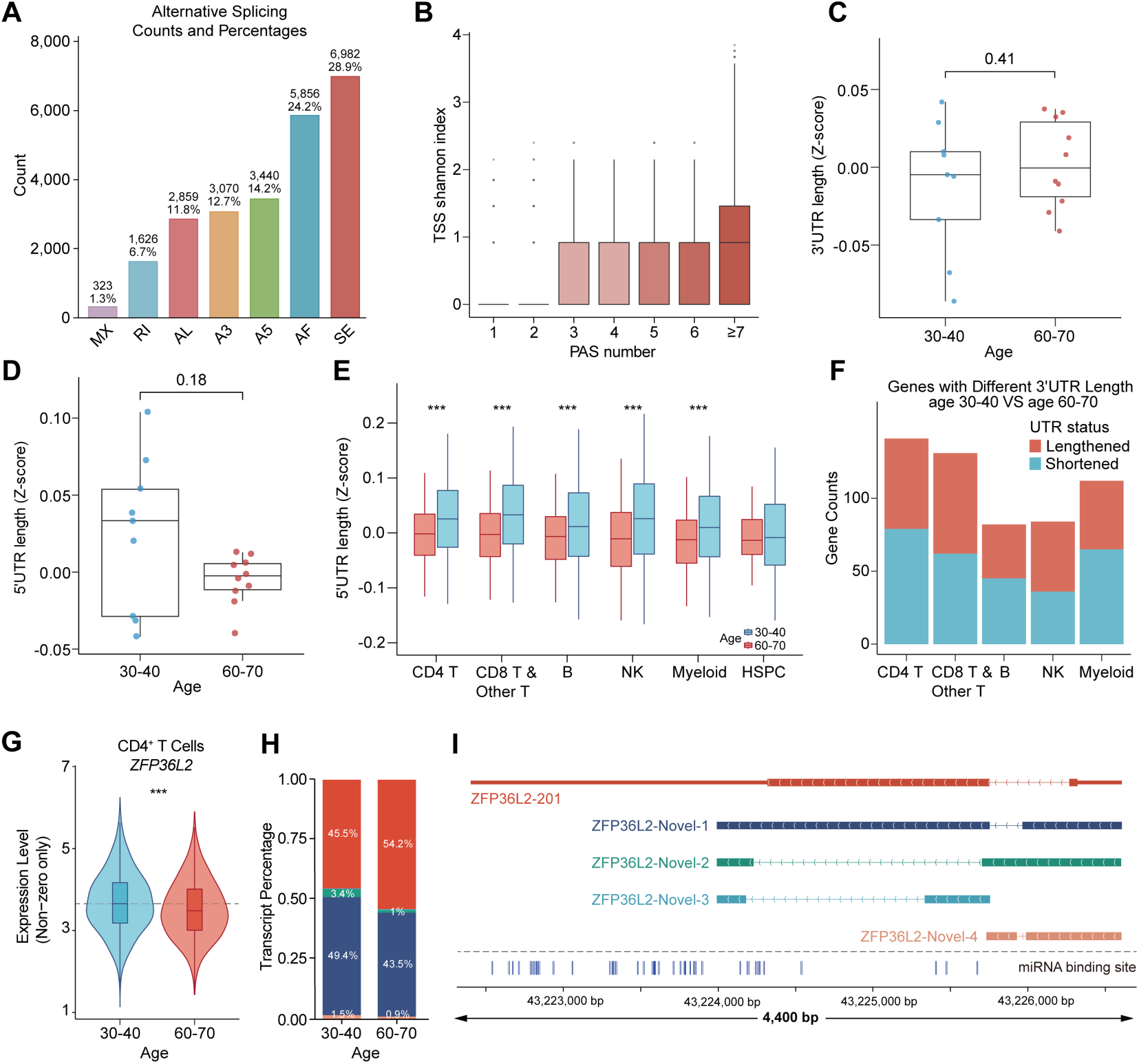
Alternative Splicing and UTR Variability Across Immune Cell Subsets with Age. **A**, Bar plot showing the number and proportion of alternative splicing (AS) events detected across all cells. **B**, 5′ end diversity, quantified using the Shannon index of all 5′-3′ isoforms, shows a positive correlation with the number of alternative polyadenylation (APA) sites per gene, indicating that genes with more 3′ ends tend to exhibit greater 5′ variability. **C-D**, Comparison of average UTR lengths between young and elderly groups. (C) Distribution of 3′ UTR lengths; (D) Distribution of 5′ UTR lengths. **E**, Boxplots of 5′-UTR lengths across immune cell subsets in the 30-40 years (blue) and 60-70 years (red) groups. **F**, Distribution of genes with 3′-UTR length differences across immune cell populations. **G**, Visualization of ZFP36L2 transcript isoforms and predicted miRNA binding sites (miRNA annotations obtained from the TargetScan database, McGeary et al., 2019). **H**, Distribution of ZFP36L2 expression across the 30-40-year (blue) and 60-70-year (red) age groups. **I**, Transcript abundance of ZFP36L2 stratified by age groups.

**Figure S11.**
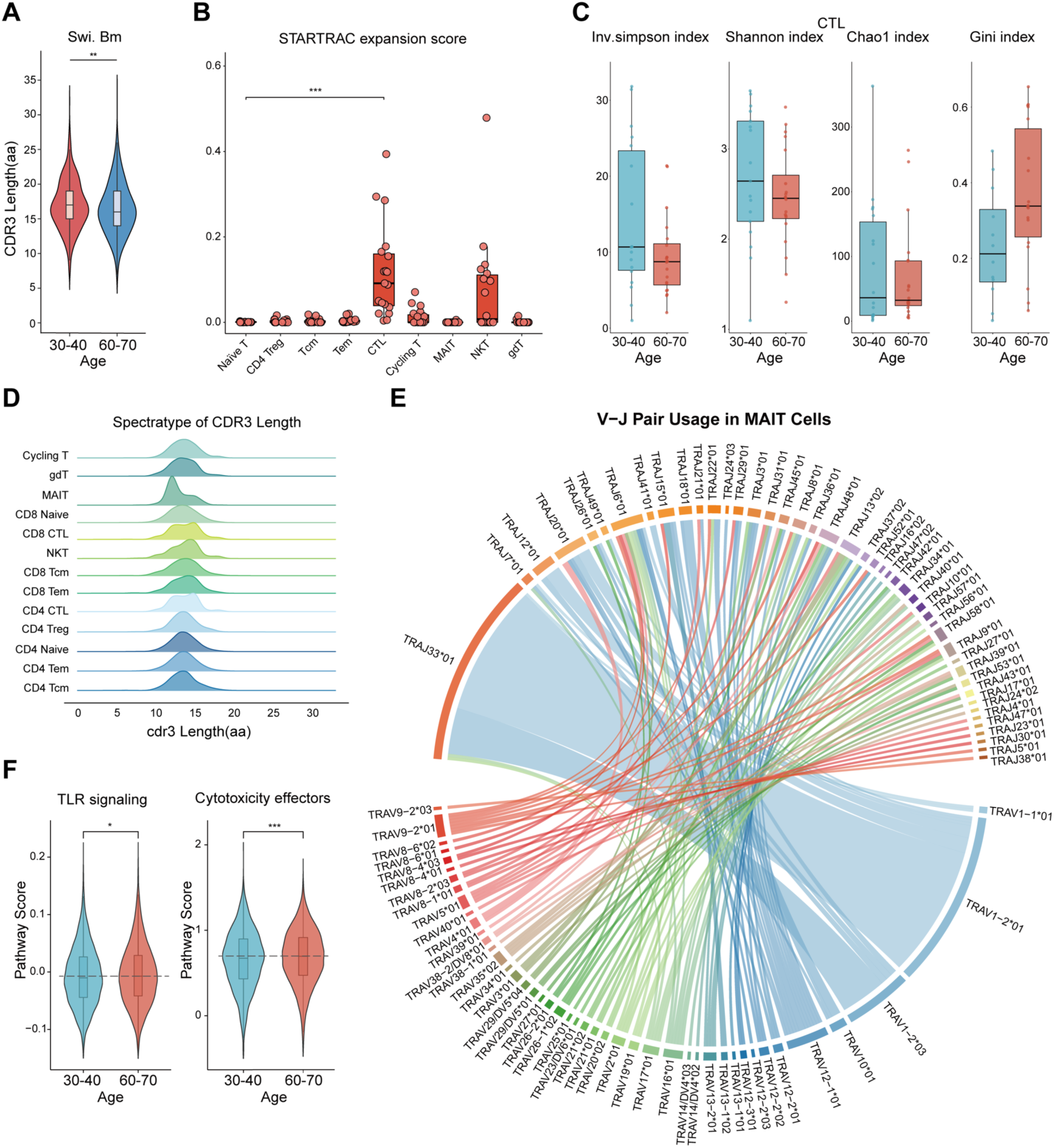
Multidimensional Quantitative Profiling of Age-Related Remodeling in the Human Immune Repertoire. **A**, IGH CDR3 length distribution in Switched Memory B. There are significant diffenrences between two groups (\*\**P* <0.01), indicating repertoire diversification (\**P* <0.05, \*\**P* <0.01, \*\*\**P* <0.001, two-sided Wilcoxon rank-sum test). **B**, Clonal expansion levels of T cell clusters quantified by STARTRAC-expansion indices for each sample (n=19). **C**, Assessment of CTL repertoire diversity across age groups using four distinct metrics (inverse Simpson, Shannon, Chao1, Gini) reveals no significant differences between individuals aged 30-40 and 60-70 years. **D**, Spectratyping of TCRα CDR3 length distribution reveals distinct repertoire profiles across human T cell subsets. **E**, Circos plot visualizing preferential V-J gene pairings in the TCR alpha chain of MAIT cells, revealing key biases in repertoire architecture. **F**, Violin plots comparing pathway scores in CTL cells between younger (30-40 years) and older (60-70 years) age groups. Statistical significance of group differences in pathway scores was determined by two-sided Wilcoxon rank-sum test.

**Figure S12.**
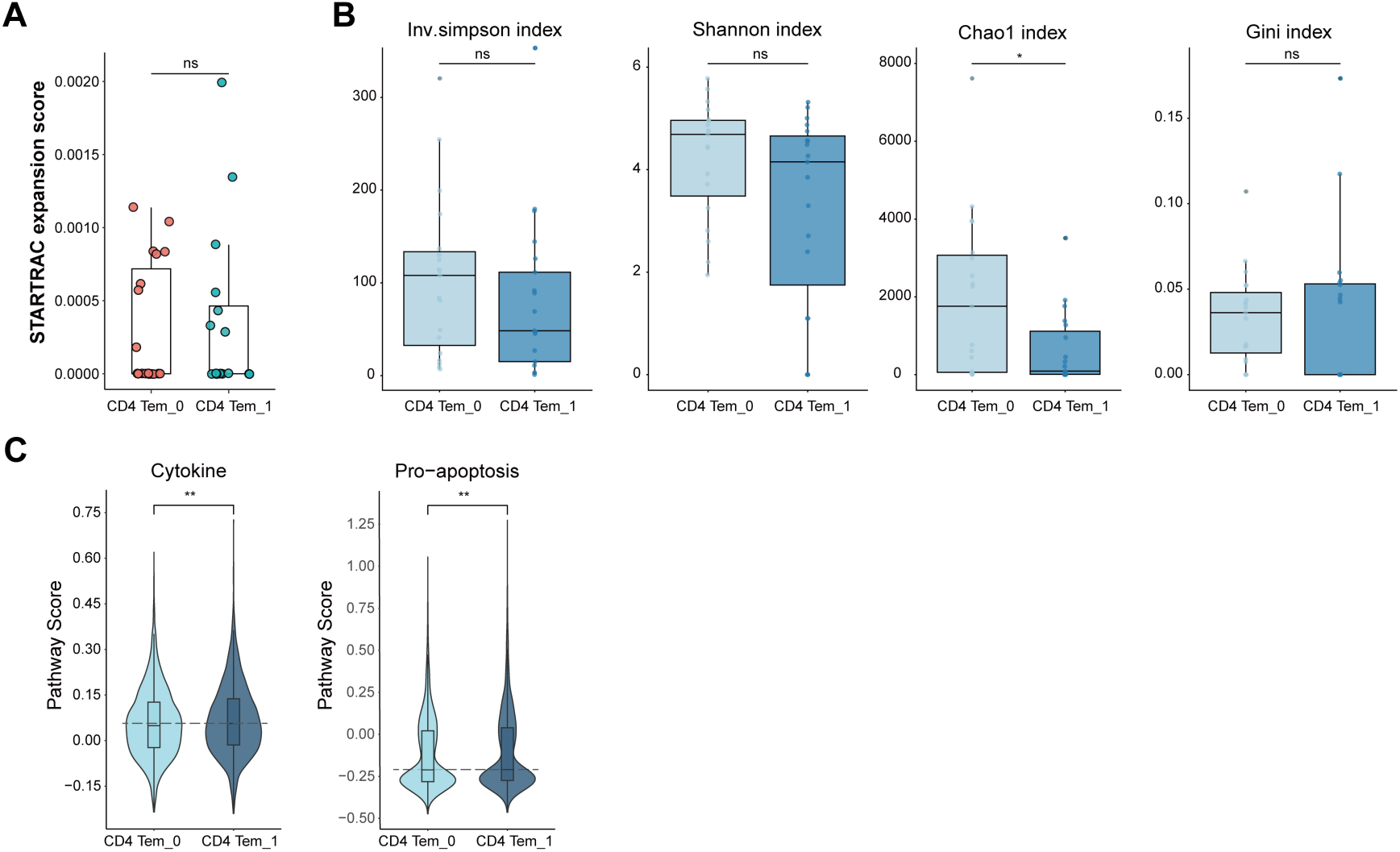
Comparative Analysis of Clonal Dynamics and Functional Programs between CD4 Tem_0 and CD4 Tem_1 Subsets. **A**, STARTRAC analysis reveals comparable clonal expansion potential between CD4 Tem_0 and CD4 Tem_1 subsets (ns, two-tailed Mann-Whitney U test). **B**, Comparative analysis of CD4 Tem_0 and CD4 Tem_1 subclusters reveals distinct clonal diversity patterns across four ecological metrics (ns, not significant; \**P* < 0.05,). **C**, Violin plots comparing pathway scores between CD4 Tem_0 and CD4 Tem_1 subclusters. Statistical significance of group differences in pathway scores was determined by two-sided Wilcoxon rank-sum test.

## References

Abugessaisa, I., Noguchi, S., Hasegawa, A., Kondo, A., Kawaji, H., Carninci, P., Kasukawa, T. (2019). refTSS: A Reference Data Set for Human and Mouse Transcription Start Sites. J Mol Biol 431, 2407–2422.

Aibar, S., Gonzalez-Blas, C.B., Moerman, T., Huynh-Thu, V.A., Imrichova, H., Hulselmans, G., Rambow, F., Marine, J., Geurts, P., Aerts, J., van den Oord, J., Atak, Z.K., Wouters, J., Aerts, S. (2017). SCENIC: single-cell regulatory network inference and clustering. Nat Methods 14, 1083–1086.

Banerjee, S., Galarza-Muñoz, G., Garcia-Blanco, M.A. (2023). Role of RNA Alternative Splicing in T Cell Function and Disease. Genes (Basel) 14, 1896.

Binnewies, M., Mujal, A.M., Pollack, J.L., Combes, A.J., Hardison, E.A., Barry, K.C., Tsui, J., Ruhland, M.K., Kersten, K., Abushawish, M.A., Spasic, M., Giurintano, J.P., Chan, V., Daud, A.I., Ha, P., Ye, C.J., Roberts, E.W., Krummel, M.F. (2019). Unleashing Type-2 Dendritic Cells to Drive Protective Antitumor CD4+ T Cell Immunity. Cell 177, 556–571.

Borcherding, N., Bormann, N.L., Kraus, G. (2020). scRepertoire: An R-based toolkit for single-cell immune receptor analysis. F1000Res 9, 47.

Borgoni, S., Kudryashova, K.S., Burka, K., de Magalhães, J.P. (2021). Targeting immune dysfunction in aging. Ageing Res Rev 70, 101410.

Britanova, O.V., Shugay, M., Merzlyak, E.M., Staroverov, D.B., Putintseva, E.V., Turchaninova, M.A., Mamedov, I.Z., Pogorelyy, M.V., Bolotin, D.A., Izraelson, M., Davydov, A.N., Egorov, E.S., Kasatskaya, S.A., Rebrikov, D.V., Lukyanov, S., Chudakov, D.M. (2016). Dynamics of Individual T Cell Repertoires: From Cord Blood to Centenarians. J Immunol 196, 5005–5013.

Chen, M., Lyu, G., Han, M., Nie, H., Shen, T., Chen, W., Niu, Y., Song, Y., Li, X., Li, H., Chen, X., Wang, Z., Xia, Z., Li, W., Tian, X., Ding, C., Gu, J., Zheng, Y., Liu, X., Hu, J., Wei, G., Tao, W., Ni, T. (2018). 3’ UTR lengthening as a novel mechanism in regulating cellular senescence. Genome Res 28, 285–294.

Chen, S. (2023). Ultrafast one-pass FASTQ data preprocessing, quality control, and deduplication using fastp. Imeta 2, e107.

Chu, Y., Dai, E., Li, Y., Han, G., Pei, G., Ingram, D.R., Thakkar, K., Qin, J., Dang, M., Le, X., Hu, C., Deng, Q., Sinjab, A., Gupta, P., Wang, R., Hao, D., Peng, F., Yan, X., Liu, Y., Song, S., Zhang, S., Heymach, J.V., Reuben, A., Elamin, Y.Y., Pizzi, M.P., Lu, Y., Lazcano, R., Hu, J., Li, M., Curran, M., Futreal, A., Maitra, A., Jazaeri, A.A., Ajani, J.A., Swanton, C., Cheng, X., Abbas, H.A., Gillison, M., Bhat, K., Lazar, A.J., Green, M., Litchfield, K., Kadara, H., Yee, C., Wang, L. (2023). Pan-cancer T cell atlas links a cellular stress response state to immunotherapy resistance. Nat Med 29, 1550–1562.

Cruz-Guilloty, F., Pipkin, M.E., Djuretic, I.M., Levanon, D., Lotem, J., Lichtenheld, M.G., Groner, Y., Rao, A. (2009). Runx3 and T-box proteins cooperate to establish the transcriptional program of effector CTLs. J Exp Med 206, 51–59.

Ferrucci, L., Fabbri, E. (2018). Inflammageing: chronic inflammation in ageing, cardiovascular disease, and frailty. Nat Rev Cardiol 15, 505–522.

Gao, S., Wu, Z., Arnold, B., Diamond, C., Batchu, S., Giudice, V., Alemu, L., Raffo, D.Q., Feng, X., Kajigaya, S., Barrett, J., Ito, S., Young, N.S. (2022). Single-cell RNA sequencing coupled to TCR profiling of large granular lymphocyte leukemia T cells. Nat Commun 13, 1982.

Garg, S.K., Delaney, C., Toubai, T., Ghosh, A., Reddy, P., Banerjee, R., Yung, R. (2014). Aging is associated with increased regulatory T-cell function. Aging Cell 13, 441–448.

Garrett Sinha, L.A. (2023). An update on the roles of transcription factor Ets1 in autoimmune diseases. WIREs Mech Dis 15, e1627.

Godec, J., Tan, Y., Liberzon, A., Tamayo, P., Bhattacharya, S., Butte, A.J., Mesirov, J.P., Haining, W.N. (2016). Compendium of Immune Signatures Identifies Conserved and Species-Specific Biology in Response to Inflammation. Immunity 44, 194–206.

Goronzy, J.J., Weyand, C.M. (2019). Mechanisms underlying T cell ageing. Nat Rev Immunol 19, 573–583.

Hamilton, S.E., Jameson, S.C. (2012). CD8 T cell quiescence revisited. Trends Immunol 33, 224–230.

Hao, Y., Hao, S., Andersen-Nissen, E., Mauck, W.M.R., Zheng, S., Butler, A., Lee, M.J., Wilk, A.J., Darby, C., Zager, M., Hoffman, P., Stoeckius, M., Papalexi, E., Mimitou, E.P., Jain, J., Srivastava, A., Stuart, T., Fleming, L.M., Yeung, B., Rogers, A.J., McElrath, J.M., Blish, C.A., Gottardo, R., Smibert, P., Satija, R. (2021). Integrated analysis of multimodal single-cell data. Cell 184, 3573–3587.

Hao, Y., Stuart, T., Kowalski, M.H., Choudhary, S., Hoffman, P., Hartman, A., Srivastava, A., Molla, G., Madad, S., Fernandez-Granda, C., Satija, R. (2024). Dictionary learning for integrative, multimodal and scalable single-cell analysis. Nat Biotechnol 42, 293–304.

Harries, L.W., Hernandez, D., Henley, W., Wood, A.R., Holly, A.C., Bradley Smith, R.M., Yaghootkar, H., Dutta, A., Murray, A., Frayling, T.M., Guralnik, J.M., Bandinelli, S., Singleton, A., Ferrucci, L., Melzer, D. (2011). Human aging is characterized by focused changes in gene expression and deregulation of alternative splicing. Aging Cell 10, 868–878.

Hetz, C. (2012). The unfolded protein response: controlling cell fate decisions under ER stress and beyond. Nat Rev Mol Cell Biol 13, 89–102.

Huang, Z., Chen, B., Liu, X., Li, H., Xie, L., Gao, Y., Duan, R., Li, Z., Zhang, J., Zheng, Y., Su, W. (2021). Effects of sex and aging on the immune cell landscape as assessed by single-cell transcriptomic analysis. Proc Natl Acad Sci U S A 118, e2023216118.

Ju, S., Zhu, Y., Liu, L., Dai, S., Li, C., Chen, E., He, Y., Zhang, X., Lu, B. (2009). Gadd45b and Gadd45g are important for anti-tumor immune responses. Eur J Immunol 39, 3010–3018.

Kennedy, B.K., Berger, S.L., Brunet, A., Campisi, J., Cuervo, A.M., Epel, E.S., Franceschi, C., Lithgow, G.J., Morimoto, R.I., Pessin, J.E., Rando, T.A., Richardson, A., Schadt, E.E., Wyss-Coray, T., Sierra, F. (2014). Geroscience: Linking Aging to Chronic Disease. Cell 159, 709–713.

Kong, S., Kim, S., Sandal, B., Lee, S., Gao, B., Zhang, D.D., Fang, D. (2011). The Type III Histone Deacetylase Sirt1 Protein Suppresses p300-mediated Histone H3 Lysine 56 Acetylation at Bclaf1 Promoter to Inhibit T Cell Activation. Journal of Biological Chemistry 286, 16967–16975.

Korsunsky, I., Millard, N., Fan, J., Slowikowski, K., Zhang, F., Wei, K., Baglaenko, Y., Brenner, M., Loh, P., Raychaudhuri, S. (2019). Fast, sensitive and accurate integration of single-cell data with Harmony. Nat Methods 16, 1289–1296.

Li, H. (2018). Minimap2: pairwise alignment for nucleotide sequences. Bioinformatics 34, 3094–3100.

Liebermann, D.A., Hoffman, B. (2014). Gadd45 in stress signaling. Journal of Molecular Signaling 3, 15.

Lin, X., Wang, X., Liu, C., Liu, C., Zeng, T., Yuan, Z., Hu, M., Xiang, R., Zhao, K., Zhou, J., Yang, S., Wang, Y., Meng, K., Wang, H., He, G., Zhao, R., Liu, J., Huang, Y., Pan, J., Wang, J., Chen, J., Guo, F., Dong, Y., Xu, X., Luo, D., Gu, Y., Liu, L., Dong, Z., Chen, L. (2024). Deciphering the Cell-Specific Transcript Heterogeneity and Alternative Splicing during the Early Embryonic Development of Zebrafish. Biorxiv.

Liu, B., Li, C., Li, Z., Wang, D., Ren, X., Zhang, Z. (2020). An entropy-based metric for assessing the purity of single cell populations. Nat Commun 11, 3155.

Liu, N., Wu, J., Deng, E., Zhong, J., Wei, B., Cai, T., Xie, Z., Duan, X., Fu, S., Osei-Hwedieh, D.O., Huang, K., Zhuang, P., Sha, O., Chen, Y., Lv, X., Zhu, Y., Zhang, L., Lin, H., Li, Q., Lu, P., Miao, J., Yamada, T., Cai, L., Du, H., Baca, S.C., Huang, Q., Ferrone, S., Wang, X., Xu, F., Fan, X., Fan, S. (2025). Immunotherapy and senolytics in head and neck squamous cell carcinoma: phase 2 trial results. Nat Med 31, 3047–3061.

Liu, Z., Liang, Q., Ren, Y., Guo, C., Ge, X., Wang, L., Cheng, Q., Luo, P., Zhang, Y., Han, X. (2023). Immunosenescence: molecular mechanisms and diseases. Signal Transduct Target Ther 8, 200.

Luo, O.J., Lei, W., Zhu, G., Ren, Z., Xu, Y., Xiao, C., Zhang, H., Cai, J., Luo, Z., Gao, L., Su, J., Tang, L., Guo, W., Su, H., Zhang, Z., Fang, E.F., Ruan, Y., Leng, S.X., Ju, Z., Lou, H., Gao, J., Peng, N., Chen, J., Bao, Z., Liu, F., Chen, G. (2022). Multidimensional single-cell analysis of human peripheral blood reveals characteristic features of the immune system landscape in aging and frailty. Nat Aging 2, 348–364.

Lynch, H.E., Goldberg, G.L., Chidgey, A., Van den Brink, M.R.M., Boyd, R., Sempowski, G.D. (2009). Thymic involution and immune reconstitution. Trends Immunol 30, 366–373.

Marcel, M. (2011). Cutadapt removes adapter sequences from high-throughput sequencing reads. EMBnet Journal 1, 10–12.

McGeary, S.E., Lin, K.S., Shi, C.Y., Pham, T.M., Bisaria, N., Kelley, G.M., Bartel, D.P. (2019). The biochemical basis of microRNA targeting efficacy. Science 366.

McGinnis, C.S., Murrow, L.M., Gartner, Z.J. (2019). DoubletFinder: Doublet Detection in Single-Cell RNA Sequencing Data Using Artificial Nearest Neighbors. Cell Syst 8, 329–337.

Mitschka, S., Mayr, C. (2022). Context-specific regulation and function of mRNA alternative polyadenylation. Nat Rev Mol Cell Biol 23, 779–796.

Mogilenko, D.A., Shpynov, O., Andhey, P.S., Arthur, L., Swain, A., Esaulova, E., Brioschi, S., Shchukina, I., Kerndl, M., Bambouskova, M., Yao, Z., Laha, A., Zaitsev, K., Burdess, S., Gillfilan, S., Stewart, S.A., Colonna, M., Artyomov, M.N. (2021a). Comprehensive Profiling of an Aging Immune System Reveals Clonal GZMK+ CD8+ T Cells as Conserved Hallmark of Inflammaging. Immunity 54, 99–115.

Mogilenko, D.A., Shpynov, O., Andhey, P.S., Arthur, L., Swain, A., Esaulova, E., Brioschi, S., Shchukina, I., Kerndl, M., Bambouskova, M., Yao, Z., Laha, A., Zaitsev, K., Burdess, S., Gillfilan, S., Stewart, S.A., Colonna, M., Artyomov, M.N. (2021b). Comprehensive Profiling of an Aging Immune System Reveals Clonal GZMK(+) CD8(+) T Cells as Conserved Hallmark of Inflammaging. Immunity 54, 99–115.

Moskalev, A.A., Smit-McBride, Z., Shaposhnikov, M.V., Plyusnina, E.N., Zhavoronkov, A., Budovsky, A., Tacutu, R., Fraifeld, V.E. (2012). Gadd45 proteins: relevance to aging, longevity and age-related pathologies. Ageing Res Rev 11, 51–66.

O’Brien, J., Hayder, H., Zayed, Y., Peng, C. (2018). Overview of MicroRNA Biogenesis, Mechanisms of Actions, and Circulation. Front Endocrinol (Lausanne) 9, 402.

Pardo-Palacios, F.J., Arzalluz-Luque, A., Kondratova, L., Salguero, P., Mestre-Tomas, J., Amorin, R., Estevan-Morio, E., Liu, T., Nanni, A., McIntyre, L., Tseng, E., Conesa, A. (2024). SQANTI3: curation of long-read transcriptomes for accurate identification of known and novel isoforms. Nat Methods 21, 793–797.

Persad, S., Choo, Z., Dien, C., Sohail, N., Masilionis, I., Chaligne, R., Nawy, T., Brown, C.C., Sharma, R., Pe’Er, I., Setty, M., Pe’Er, D. (2023). SEACells infers transcriptional and epigenomic cellular states from single-cell genomics data. Nat Biotechnol 41, 1746–1757.

Piccirillo, C.A., Bjur, E., Topisirovic, I., Sonenberg, N., Larsson, O. (2014). Translational control of immune responses: from transcripts to translatomes. Nat Immunol 15, 503–511.

Porcelli, S., Yockey, C.E., Brenner, M.B., Balk, S.P. (1993). Analysis of T cell antigen receptor (TCR) expression by human peripheral blood CD4-8-alpha/beta T cells demonstrates preferential use of several V beta genes and an invariant TCR alpha chain. J Exp Med 178, 1–16.

Prjibelski, A.D., Mikheenko, A., Joglekar, A., Smetanin, A., Jarroux, J., Lapidus, A.L., Tilgner, H.U. (2023). Accurate isoform discovery with IsoQuant using long reads. Nat Biotechnol 41, 915–918.

Ramos Pittol, J.M., Oruba, A., Mittler, G., Saccani, S., van Essen, D. (2018). Zbtb7a is a transducer for the control of promoter accessibility by NF-kappa B and multiple other transcription factors. PLoS Biol 16, e2004526.

Redmond, D., Poran, A., Elemento, O. (2016). Single-cell TCRseq: paired recovery of entire T-cell alpha and beta chain transcripts in T-cell receptors from single-cell RNAseq. Genome Med 8, 80.

Rodríguez-Jiménez, P., Fernández-Messina, L., Ovejero-Benito, M.C., Chicharro, P., Vera-Tomé, P., Vara, A., Cibrian, D., Martínez-Fleta, P., Jiménez-Fernández, M., Sánchez-García, I., Llamas-Velasco, M., Abad-Santos, F., Sánchez-Madrid, F., Dauden, E., de la Fuente, H. (2021). Growth arrest and DNA damage-inducible proteins (GADD45) in psoriasis. Sci Rep 11, 14579.

Satija, R., Farrell, J.A., Gennert, D., Schier, A.F., Regev, A. (2015). Spatial reconstruction of single-cell gene expression data. Nat Biotechnol 33, 495–502.

Sauce, D., Larsen, M., Fastenackels, S., Duperrier, A., Keller, M., Grubeck-Loebenstein, B., Ferrand, C., Debré, P., Sidi, D., Appay, V. (2009). Evidence of premature immune aging in patients thymectomized during early childhood. J Clin Invest 119, 3070–3078.

Shen, W., Le, S., Li, Y., Hu, F. (2016). SeqKit: A Cross-Platform and Ultrafast Toolkit for FASTA/Q File Manipulation. PLoS One 11, e0163962.

Smith, T., Heger, A., Sudbery, I. (2017). UMI-tools: modeling sequencing errors in Unique Molecular Identifiers to improve quantification accuracy. Genome Res 27, 491–499.

Song, C., Pan, W., Brown, B., Tang, C., Huang, Y., Chen, H., Peng, N., Wang, Z., Weber, D., Byrne-Steele, M., Wu, H., Liu, H., Deng, Y., He, N., Li, S. (2022). Immune repertoire analysis of normal Chinese donors at different ages. Cell Prolif 55, e13311.

Stahl, E.C., Brown, B.N. (2015). Cell Therapy Strategies to Combat Immunosenescence. Organogenesis 11, 159–172.

Stephensen, C.B., Borowsky, A.D., Lloyd, K.C.K. (2007). Disruption of*Rxra* gene in thymocytes and T lymphocytes modestly alters lymphocyte frequencies, proliferation, survival and T helper type 1/type 2 balance. Immunology 121, 484–498.

Sturm, G., Szabo, T., Fotakis, G., Haider, M., Rieder, D., Trajanoski, Z., Finotello, F. (2020). Scirpy: a Scanpy extension for analyzing single-cell T-cell receptor-sequencing data. Bioinformatics 36, 4817–4818.

Suo, C., Polanski, K., Dann, E., Lindeboom, R.G.H., Vilarrasa-Blasi, R., Vento-Tormo, R., Haniffa, M., Meyer, K.B., Dratva, L.M., Tuong, Z.K., Clatworthy, M.R., Teichmann, S.A. (2024). Dandelion uses the single-cell adaptive immune receptor repertoire to explore lymphocyte developmental origins. Nat Biotechnol 42, 40–51.

Ten Broeke, T., Wubbolts, R., Stoorvogel, W. (2013). MHC Class II Antigen Presentation by Dendritic Cells Regulated through Endosomal Sorting. Cold Spring Harb Perspect Biol 5, a016873.

Terekhova, M., Swain, A., Bohacova, P., Aladyeva, E., Arthur, L., Laha, A., Mogilenko, D.A., Burdess, S., Sukhov, V., Kleverov, D., Echalar, B., Tsurinov, P., Chernyatchik, R., Husarcikova, K., Artyomov, M.N. (2023a). Single-cell atlas of healthy human blood unveils age-related loss of NKG2C(+)GZMB(-)CD8(+) memory T cells and accumulation of type 2 memory T cells. Immunity 56, 2836–2854.

Terekhova, M., Swain, A., Bohacova, P., Aladyeva, E., Arthur, L., Laha, A., Mogilenko, D.A., Burdess, S., Sukhov, V., Kleverov, D., Echalar, B., Tsurinov, P., Chernyatchik, R., Husarcikova, K., Artyomov, M.N. (2023b). Single-cell atlas of healthy human blood unveils age-related loss of NKG2C(+)GZMB(-)CD8(+) memory T cells and accumulation of type 2 memory T cells. Immunity 56, 2836–2854.

Terekhova, M., Swain, A., Bohacova, P., Aladyeva, E., Arthur, L., Laha, A., Mogilenko, D.A., Burdess, S., Sukhov, V., Kleverov, D., Echalar, B., Tsurinov, P., Chernyatchik, R., Husarcikova, K., Artyomov, M.N. (2024). Single-cell atlas of healthy human blood unveils age-related loss of NKG2C(+)GZMB(-)CD8(+) memory T cells and accumulation of type 2 memory T cells. Immunity 57, 188–192.

Thaxton, J.E., Wallace, C., Riesenberg, B., Zhang, Y., Paulos, C.M., Beeson, C.C., Liu, B., Li, Z. (2017). Modulation of Endoplasmic Reticulum Stress Controls CD4+ T-cell Activation and Antitumor Function. Cancer Immunol Res 5, 666–675.

Tilloy, F., Treiner, E., Park, S.H., Garcia, C., Lemonnier, F., de la Salle, H., Bendelac, A., Bonneville, M., Lantz, O. (1999). An invariant T cell receptor alpha chain defines a novel TAP-independent major histocompatibility complex class Ib-restricted alpha/beta T cell subpopulation in mammals. J Exp Med 189, 1907–1921.

Treiner, E., Duban, L., Bahram, S., Radosavljevic, M., Wanner, V., Tilloy, F., Affaticati, P., Gilfillan, S., Lantz, O. (2003). Selection of evolutionarily conserved mucosal-associated invariant T cells by MR1. Nature 422, 164–169.

Vitting-Seerup, K., Sandelin, A. (2019). IsoformSwitchAnalyzeR: analysis of changes in genome-wide patterns of alternative splicing and its functional consequences. Bioinformatics 35, 4469–4471.

Walford, R.L. (1964). The Immunologic Theory of Aging. Gerontologist 4, 195–197.

Wang, Y., Li, R., Tong, R., Chen, T., Sun, M., Luo, L., Li, Z., Chen, Y., Zhao, Y., Zhang, C., Wei, L., Lin, W., Chen, H., Qian, K., Chen, A.F., Liu, J., Chen, L., Li, B., Wang, F., Wang, L., Su, B., Pu, J. (2025a). Integrating single-cell RNA and T cell/B cell receptor sequencing with mass cytometry reveals dynamic trajectories of human peripheral immune cells from birth to old age. Nat Immunol 26, 308–322.

Wang, Y., Li, R., Tong, R., Chen, T., Sun, M., Luo, L., Li, Z., Chen, Y., Zhao, Y., Zhang, C., Wei, L., Lin, W., Chen, H., Qian, K., Chen, A.F., Liu, J., Chen, L., Li, B., Wang, F., Wang, L., Su, B., Pu, J. (2025b). Integrating single-cell RNA and T cell/B cell receptor sequencing with mass cytometry reveals dynamic trajectories of human peripheral immune cells from birth to old age. Nat Immunol 26, 308–322.

Wu, T., Hu, E., Xu, S., Chen, M., Guo, P., Dai, Z., Feng, T., Zhou, L., Tang, W., Zhan, L., Fu, X., Liu, S., Bo, X., Yu, G. (2021). clusterProfiler 4.0: A universal enrichment tool for interpreting omics data. Innovation (Camb) 2, 100141.

Ye, J., Huang, X., Hsueh, E.C., Zhang, Q., Ma, C., Zhang, Y., Varvares, M.A., Hoft, D.F., Peng, G. (2012). Human regulatory T cells induce T-lymphocyte senescence. Blood 120, 2021–2031.

Yin, J., Zheng, Y., Huang, Z., Zhou, W., Yuan, Y., Cai, P., Bai, Y., Yang, S., Gao, Y., Duan, S., Wang, Y., Zhang, W., Zhang, X., Wei, Y., Xu, Z., Huang, Y., Liu, Y., Wang, W., Yang, T., Lv, J., Zhang, Z., Chen, X., Zhang, X., Li, F., Zhang, Y., Zeng, G., Wang, X., Ma, W., Hou, G., Hao, S., Liu, C., Lai, Y., Wang, B., Li, Y., Zhang, W., Gao, P., Xie, J., Esteban, M.A., Gu, Y., Ji, J., Qi, T., Liu, B., Wang, J., Yang, J., Xu, X., Liu, L., Jin, X., Liu, C. (2025). Single-Cell Genomics Elucidates Molecular Variations and Regulatory Mechanisms in Circulating Immune Cells. Biorxiv.

Yong, C., Liang, Y., Wang, M., Jin, W., Fan, X., Wang, Z., Cao, K., Wu, T., Li, Q., Chang, C. (2025). Alternative splicing: A key regulator in T cell response and cancer immunotherapy. Pharmacol Res 215, 107713.

Zhang, L., Yu, X., Zheng, L., Zhang, Y., Li, Y., Fang, Q., Gao, R., Kang, B., Zhang, Q., Huang, J.Y., Konno, H., Guo, X., Ye, Y., Gao, S., Wang, S., Hu, X., Ren, X., Shen, Z., Ouyang, W., Zhang, Z. (2018). Lineage tracking reveals dynamic relationships of T cells in colorectal cancer. Nature 564, 268–272.

Zhong, J., Ding, R., Jiang, H., Li, L., Wan, J., Feng, X., Chen, M., Peng, L., Li, X., Lin, J., Yang, H., Wang, M., Li, Q., Chen, Q. (2023). Single-cell RNA sequencing reveals the molecular features of peripheral blood immune cells in children, adults and centenarians. Front Immunol 13, 1081889.

Zhou, P., Shi, H., Huang, H., Sun, X., Yuan, S., Chapman, N.M., Connelly, J.P., Lim, S.A., Saravia, J., Kc, A., Pruett-Miller, S.M., Chi, H. (2023). Single-cell CRISPR screens in vivo map T cell fate regulomes in cancer. Nature 624, 154–163.

